# Reversible therapeutic resistance in *EGFR* mutant lung cancer caused by *RB1* loss induced lineage plasticity

**DOI:** 10.1101/2025.08.04.668485

**Authors:** Letian Zhang, Meenalakshmi Chinnam, Bradley Balk, Xiaojing Zhang, Yanqing Wang, Tatiana Shaurova, Shamshad Alam, Saraswati Pokharel, Prashant Singh, Justine Jacobi, Jianmin Wang, Eduardo Cortes Gomez, Michael Habitzruther, Mukund Seshadri, Boyko Atanassov, Wei Wu, Elizaveta V. Benevolenskaya, Collin M. Blakely, Pamela Hershberger, Mark Long, Trever Bivona, David W. Goodrich

**Author notes:** Department of Pathology, National Cancer Center, National Clinical Research Center for Cancer, Cancer Hospital, Chinese Academy of Medical Sciences and Peking Union Medical College, Beijing, People’s Republic of China (Letian Zhang).

## Abstract

Treatment with EGFR tyrosine kinase inhibitors (EGFR TKI) is standard of care for patients with lung cancer initiated by activating mutations in the *EGFR* gene. While EGFR TKI treatment is effective, virtually all patients progress on therapy. Loss of function mutations in the *RB1* tumor suppressor gene are associated with poor EGFR TKI outcomes, but underlying mechanisms remain unclear. Analysis of genetically engineered experimental models and human clinical specimens described here demonstrates *RB1* loss has minimal impact on initial cell cycle responses to EGFR TKI but markedly accelerates acquired resistance over time. *RB1* loss promotes transcriptional plasticity that enables rare cells to spontaneously assume reversible, resistance-conferring lineage states. These findings highlight the importance of non-genetic adaptations in driving acquired EGFR TKI resistance, advance our understanding of how *RB1* impacts cancer, and identify opportunities to improve EGFR TKI treatment outcomes by targeting reversible epigenetic resistance mechanisms.

## Introduction

Lung cancer is the leading cause of cancer-mediated mortality worldwide, but patient outcomes have improved considerably with the clinical deployment of molecularly targeted therapies. EGFR tyrosine kinase inhibitors (EGFR TKI), for example, have become standard of care for lung adenocarcinoma (LUAD) patients whose disease is caused by activating mutations in the *EGFR* gene ^1–5^. Despite beneficial clinical responses, EGFR TKI are not curative as virtually all patients eventually progress on therapy. Acquired EGFR TKI resistance is often caused by secondary mutations in *EGFR* that reduce relative drug binding^6,7^. Acquired resistance can also be caused by off-target genetic mechanisms like *MET* gene amplification that activates compensatory signaling pathways^8^. Such genetic resistance mechanisms have been addressed clinically by deployment of mutation selective EGFR TKI like Osimertinib or by drugs like Crizotinib that inhibit compensatory signaling pathways. Despite extensive DNA sequencing efforts, however, genetic mechanisms driving acquired EGFR TKI resistance have not been identified in about half of all patients examined ^9–11^. Therapeutic options remain limited for such patients.

EGFR mutant LUAD lacking a functional *RB1* tumor suppressor gene is associated with inferior clinical outcomes on EGFR TKI^12–16^. Several *RB1*-regulated mechanisms could potentially account for these observations. RB1 protein is a well-established negative regulator of the cell cycle^17^, and RB1 loss could abrogate cell cycle arrest induced by EGFR TKI^18^. RB1 loss also compromises genome stability^19,20^, and rapid accumulation of resistance conferring genetic alterations in the absence of *RB1* may accelerate acquired EGFR TKI resistance. *RB1* loss of function mutations are highly recurrent in EGFR mutant LUAD that undergoes histological transformation to small cell lung cancer-like variants (tSCLC) during EGFR TKI therapy^12,13,21–24^, suggesting a role for *RB1* in lung cancer lineage plasticity. However, most *RB1* deficient LUAD do not undergo tSCLC transformation yet still exhibit comparatively poor responses to EGFR TKI treatment^12,13,22^. The mechanisms by which *RB1* loss of function drives poor EGFR TKI outcomes thus remain unknown for most affected patients. In experimental models, dynamic molecular fluctuations prime some cancer cells to survive targeted therapies, including EGFR TKI, as quiescent drug tolerant persistors (DTP)^25–32^. While most DTP remain arrested in the presence of drug, rare DTP can resume cell proliferation to seed acquired therapeutic resistance. Dynamic transcriptional changes have also been observed in clinical specimens isolated from patients at different stages of response and progression through EGFR TKI treatment^33^. These dynamic transcriptional adaptations facilitate LUAD cell survival and acquired EGFR TKI resistance. Little is known about the pathways that regulate transcriptional plasticity itself, however, and thus control the rate at which LUAD cells adapt and progress through EGFR TKI therapy.

Experiments described here use newly developed experimental models and unique clinical specimens to investigate how *RB1* loss of function affects acquired EGFR TKI resistance in LUAD. *RB1* loss has minimal impact on the cancer cell cycle, genome instability, or initial transcriptional responses to EGFR TKI in vitro, but markedly increases the rate at which LUAD cells progress through EGFR TKI therapy over time. Accelerated acquired EGFR TKI resistance is caused by *RB1*-loss induced transcriptional plasticity that enables rare treatment naïve LUAD cells to assume reversible, EGFR TKI resistant lineage states. Analysis of genetically engineered mice indicate *RB1/TP53* loss facilitates escape from dependence on mutant EGFR signaling in vivo via lineage plasticity. These observations demonstrate that alterations in genes that control epigenetic fidelity, like *RB1*, can drive acquired EGFR TKI resistance indirectly by enabling resistance conferring transcriptional adaptations. These findings highlight the importance of reversible non-genetic adaptations in acquired EGFR TKI resistance, suggest new opportunities for improving EGFR TKI therapeutic outcomes by targeting such adaptations, and advance understanding of how the *RB1* tumor suppressor gene impacts cancer.

## Results

### RB1 deficiency accelerates progression through EGFR TKI in vitro

We stably suppressed *RB1* protein (Rb) expression in three established human LUAD cell lines with activating *EGFR* mutations (**Figure S1A-D**). All cell lines were originally *RB1* wild type, had *TP53* mutations, and contained different activating EGFR mutations (H1975=L858R/T790M; HCC827=ΔEx19; PC9=ΔEx19). While *RB1* was wild type, each cell line contained genetic alterations in upstream *RB1* pathway genes that can disrupt normal Rb-mediated cell cycle control including *CDKN2A* loss (H1975, PC9), *CCND1* amplification (H1975), or *CDK4/6* amplification (H1975, HCC827)^34,35^. Suppressing or deleting *RB1* did not cause consistent changes in cell growth rate or cell cycle phase distribution in these lung cancer cell lines (**Figure 1A-B**). Nor did *RB1* deficiency consistently affect sensitivity of the cell population to the EGFR TKI Osimertinib (OSM) in 3-day dose response experiments (**Figure 1C**). The 50% inhibitory concentration (IC50) was unchanged for two cell lines after *RB1* loss (HCC827, PC9), consistent with prior reports^13^, but showed a modest increase in H1975 cells.

**Figure 1:**
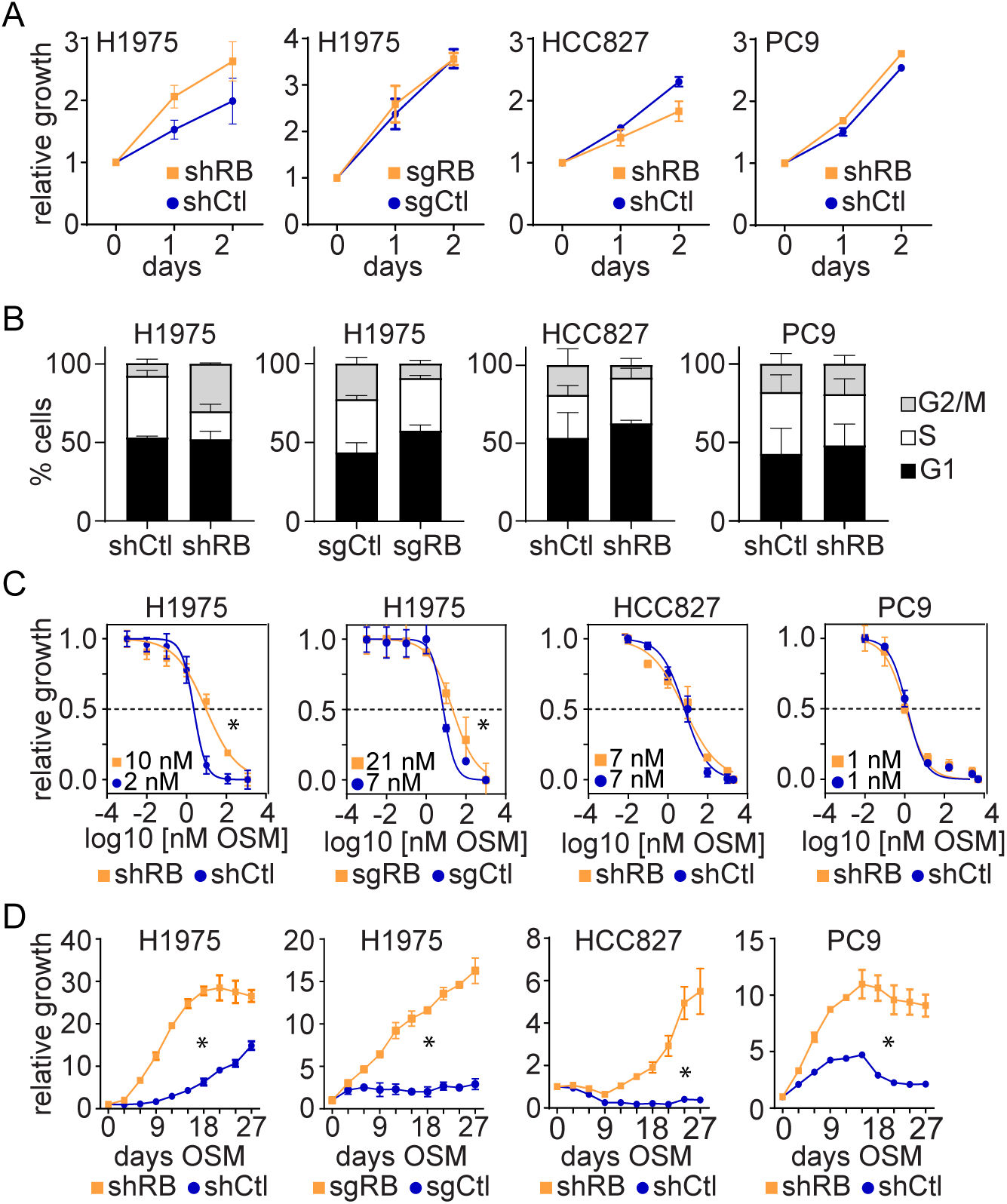
RB1 deficiency accelerates acquired EGFR TKI resistance in human LUAD cell lines. A) The listed cell lines were modified as indicated and accumulation of viable cells was measured over time in three replicate experiments using standard culture conditions. The growth of control versus *RB1* modified cells was not significantly different by 2-way ANOVA (P>0.05). B) The cell cycle phase distribution of cells in A) was measured by propidium iodide staining and flow cytometry and plotted as percent of total cells. C) The sensitivity of modified cells in A) were measured in 3-day OSM dose response experiments and the non-linear least squares fit of the data plotted. The best-fit IC50 concentrations are shown. Asterisks indicate curves were significantly different P<0.05). D) The indicated cells were exposed to OSM (1 µM) continuously for several weeks and cell accumulation monitored over time. Asterisks indicate significant differences between control and RB1 modified cells (P<0.05 two-way ANOVA).

To experimentally model time to progression on therapy, cells were treated continuously with a single clinically relevant OSM concentration (1 μM) and cell accumulation measured over several weeks. Both control and *RB1* deficient cells were sensitive to OSM, but some cells survived as DTP as expected. After prolonged OSM exposure, some DTP of both genotypes resumed proliferation in the presence of OSM to generate expanding drug tolerant persisters (DTEP)^30,36,37^. However, *RB1* deficient cells exhibited a significantly shorter time to DTEP progression than control cells for all cell lines examined (**Figure 1D**). Similar results were obtained with an earlier generation EGFR TKI, erlotinib, indicating effects were not specific to OSM (**Figure S1E-F**). Single cell clones from both control and *RB1* deficient H1975 cells were isolated, expanded, and analyzed as above to reduce pre-existing genetic heterogeneity. The growth rates of individual clones varied, but the average doubling times of *RB1* deficient and control clones were not significantly different (**Figure S1G**). *RB1* deficient clones exhibited a shorter time to OSM progression than control clones, excepting one control clone (C5) that exhibited a time to progression comparable to *RB1* deficient clones (**Figure S1H**). *RB1* loss thus failed to cause consistent changes in the cell cycle, cell proliferation, or OSM sensitivity but uniformly accelerated the time to progression on OSM.

Cells were monitored by time lapse video microscopy to assess the heterogeneity of OSM responses at the level of individual cells. After 1-2 days of OSM exposure, both *RB1* deficient and control cells exited the cell cycle as the fraction of cells undergoing cell division in 8-hour time windows declined to undetectable levels (**Figure 2A**). *RB1* loss, therefore, did not compromise initial cell cycle arrest induced by OSM. In control cells, cell division events remained low or undetectable throughout the remainder of the 9-day experiment. In contrast, rare *RB1* deficient cells underwent sporadic cell division events beginning 96 hours after initial OSM exposure. Similar results were observed in HCC827 cells, although these cells responded more rapidly and completely to OSM (**Figure 2B**). These sporadic cell division events increased the accumulation of viable Rb deficient cells relative to controls over time (**Figure S2A-B**), but only rare cell division events gave rise to continuously proliferating DTEP colonies (**Figure 2C-D**). Across multiple experimental replicates, 19 DTEP colonies were detected within 9 days from 95,000 *RB1* deficient H1975 cells (0.02%) and 13 DTEP colonies were detected from 107,000 RB1 deficient HCC827 cells (0.012%). No DTEP colonies were detected within 9 days for control cells. Colony formation assays conducted over multiple weeks gave similar results. The colony forming potential of Rb deficient cells in the presence of OSM was significantly greater than control cells (9-fold H1975, 10-fold HCC827), even allowing control cells more time to expand into colonies (**Figure 2E-F**). Expanded single cell clones gave analogous results suggesting pre-existing genetic heterogeneity did not account for observed differences (**Figure S2C-D**). These findings indicated that a small fraction of *RB1* deficient cells regained proliferative potential during continuous OSM exposure much sooner than control cells.

**Figure 2:**
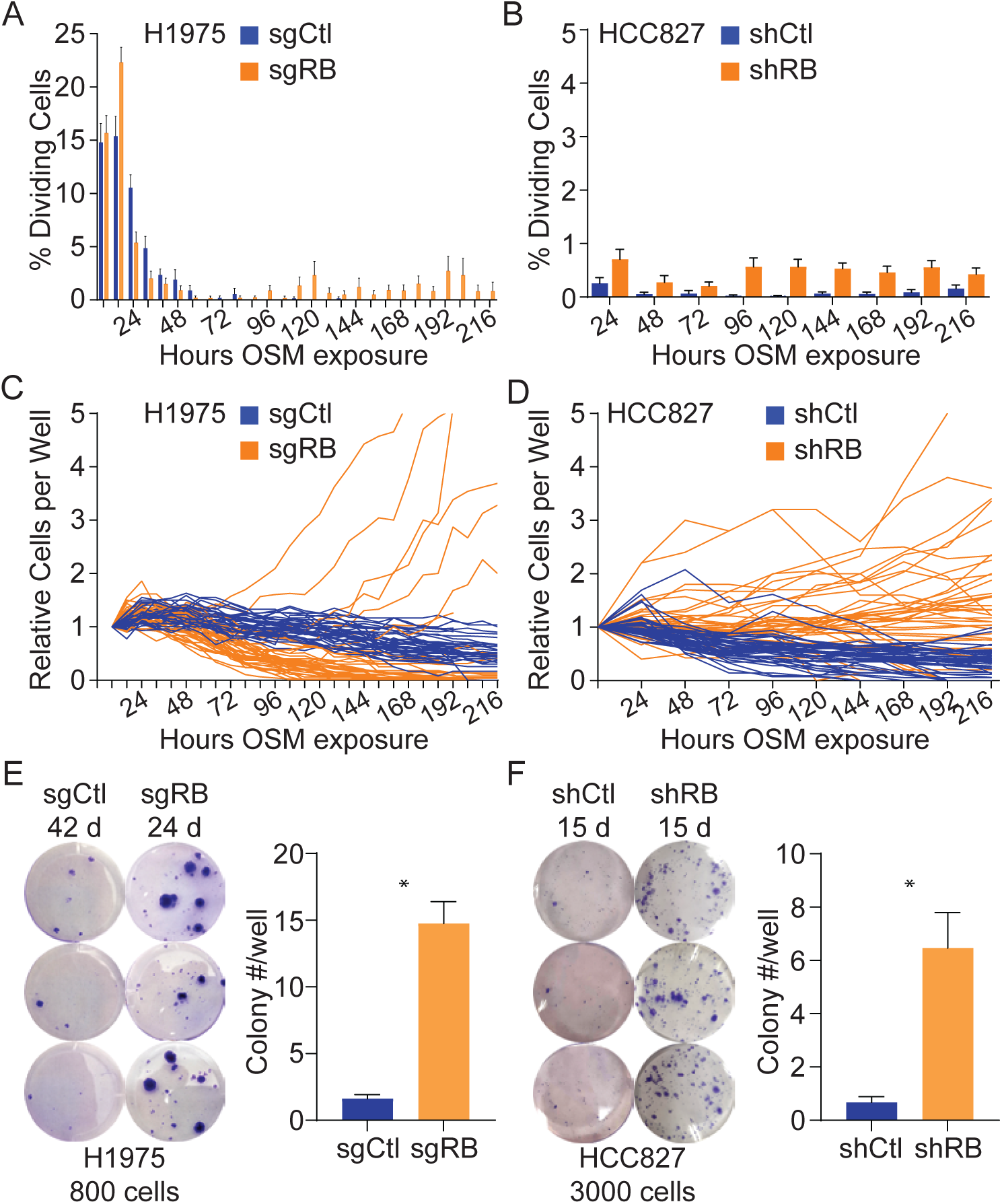
Rare *RB1* deficient LUAD cells rapidly progress to OSM DTEP. A) and B) The indicated cells were plated and treated with 1 µM OSM continuously for 9 days and the cell cycle response of individual cells was monitored by time-lapse video microscopy. The fraction of cells undergoing cell division within 8-hour time windows was calculated by manual inspection. C) and D) The indicated cells were plated in multi-well plates, treated with 1 µM OSM continuously, and accumulation of viable cells monitored over nine days. Each line represent viable cells over time in an individual well. E) and F) The listed cells were plated in 6-well plates and exposed to 1 µM OSM continuously for the indicated number of days. Cells were then fixed and stained to quantify OSM resistant colony formation. Images at left show results from a representative replicate, graphs at right show the mean and standard error of OSM resistant colonies from four or more biological replicates. Asterisks indicate statistically significant differences (T-test P<0.05).

### Known resistance conferring genetic alterations are not detected in OSM resistant cells

To test if *RB1* deficiency drives acquired OSM resistance by increasing genome instability and the acquisition of resistance conferring genetic alterations, genomic DNA was analyzed by DNA sequencing. Targeted DNA sequencing of *EGFR* exons 19-21 indicated H1975 and HCC827 OSM DTEP, both control and *RB1* deficient, retained their original *EGFR* mutations but did not acquire additional *EGFR* mutations in the exons analyzed (**Table S1**). Further, EGFR activation, as assessed by EGFR protein phosphorylation, was not detectable in OSM DTEP compared to treatment naïve (TN) cells (**Figure 3A**). Thus OSM was hitting its intended target, precluding the possibility *RB1* loss altered OSM stability or activity. Instead, activation of EGFR was not required for downstream activation of MAPK and AKT growth signaling in OSM DTEP cells. Thus off-target mechanisms were likely driving progression through OSM under the conditions used. This contrasts with prior studies that detected frequent resistance conferring second site *EGFR* mutations after deploying a dose escalation strategy for isolating EGFR TKI resistant cells^38^.

**Figure 3:**
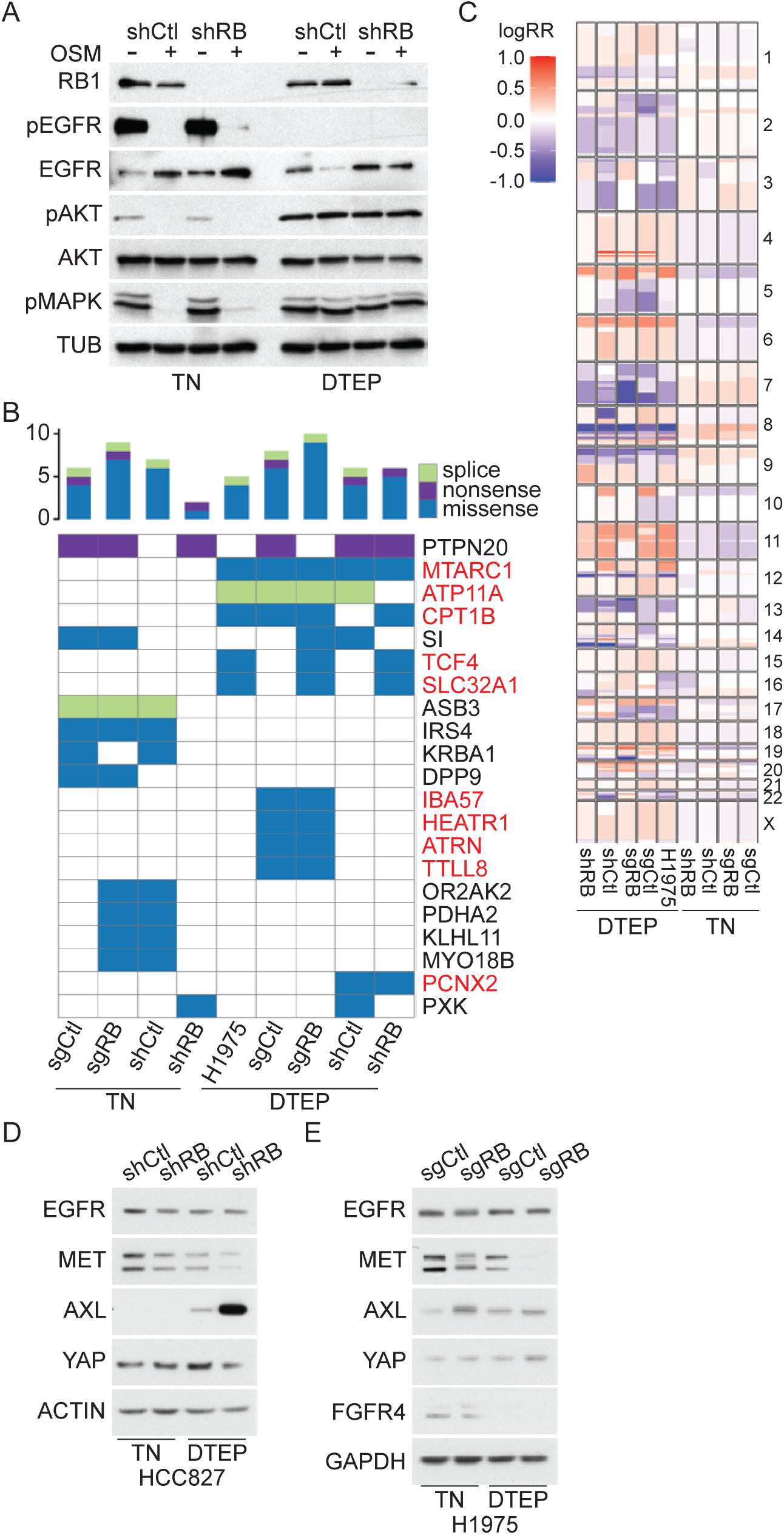
Genetic mutations do not account for accelerated OSM resistance in *RB1* deficient LUAD cells. A) Protein was extracted from treatment naïve (TN) or OSM resistant (DTEP) H1975 cells modified with the indicated shRNA, either in the presence or absence of OSM, and extracts analyzed by western blotting for the indicated proteins. Tubulin serves as a protein loading control. DTEP cell lines were established by culturing in the presence of OSM for >40 days. B) DNA was extracted from TN or OSM DTEP H1975 cells modified as indicated and analyzed by whole exome DNA sequencing. Genes with protein coding mutations detected in more than one sample, relative to the untreated parental H1975 cell reference genome, are listed. Those mutations unique to DTEP are listed in red. C) DNA copy number changes were inferred from the DNA sequencing data and plotted across chromosomes for each sample, color coded by gain (red) or loss (blue). D) Protein extracts from the indicated HCC827 samples were analyzed for the indicated proteins by western blotting. Actin serves as the protein loading control. E) Protein extracts from the indicated H1975 samples were analyzed for the indicated proteins by western blotting. GAPDH serves as the protein loading control.

Whole exome DNA sequencing was performed to test if OSM DTEP acquired known off-target resistance conferring genetic alterations (**Table S2**)^39^. Overall, the mean number of single nucleotide variants or small deletions/insertions detected in *RB1* deficient cells at baseline was not significantly different than control cells (35 versus 36, T-test P=0.97). The mean number of such DNA alterations was higher in OSM DTEP cells relative to their TN counterparts (50 versus 21, T-test P=0.018). We identified 10 genes that had recurrent protein coding mutations in more than one OSM DTEP sample (**Figure 3B**). The specific mutations detected within each recurrence were identical, suggesting they were not acquired independently and likely reflect selection of rare pre-existing sub-clones. None of the recurrent mutations unique to OSM DTEP have been associated with OSM resistance, to our knowledge, but they could provide a fitness advantage during OSM treatment. Nonetheless, none of the mutations detected were unique to *RB1* deficient cells, so they did not explain observed accelerated acquired OSM resistance.

Copy number variations were inferred from DNA sequencing data to test whether acquired OSM resistance was associated with known resistance conferring gene amplifications or deletions^39^. OSM DTEP had more copy number changes than TN cells (**Figure 3C**)(Wilcox rank test P=0.019). Twenty-one focal deletion (n=13) or amplification (n=8) peaks unique to OSM DTEP were identified with copy number changes ranging from 2.7 to −1.3. (**Figure S3, Table S3-4**). The mean copy number change at these regions was not significantly different between *RB1* deficient and control cells (−0.21 versus −0.02, T-test P=0.33). Established EGFR TKI resistance conferring copy number alterations like *MET*, *HER2*, or *EGFR* gene amplifications were not detected, nor was increased *EGFR* or *MET* protein expression detected (**Figure 3D,E**). One focal amplification was detected at 8q24.21 that spanned 16 protein coding genes, including *MYC*. Increased *MYC* protein expression has been implicated in OSM resistance^39–41^. However, this region was amplified in both *RB1* deficient and control OSM DTEP cells, 2.4- and 2.5-fold respectively, indicating this copy number change did not account for accelerated OSM progression upon *RB1* loss.

RNA-seq analysis of parental and OSM resistant cells was performed to detect potential gene fusions that might drive OSM resistance like those involving *ALK*, *BRAF*, or *RET* ^39^. *RB1* loss did not affect the number of gene fusions detected in HCC827 or H1975 cells, TN or OSM DTEP (**Tables S5-6**). H1975 OSM DTEP cells had more detectable gene fusions than TN cells (14.3 vs. 4, T test P<0.01), but OSM resistant HCC827 cells had fewer (10.2 vs. 18.8, T test P<0.01). As with somatic nucleotide variants, the recurrent gene fusions detected in multiple samples were identical, suggesting they reflected pre-existing sub-clonal genetic variants. We did not detect gene fusions known to drive EGFR TKI resistance, nor were any gene fusions uniquely recurrent in *RB1* deficient cells. Overall these results indicated *RB1* loss did not have a significant impact on genome instability over the time frames studied, and genetic alterations were unlikely to mediate effects on acquired OSM resistance.

### RB1 deficiency drives transcriptional diversity and increases the fraction of cells assuming OSM resistant transcriptional states

Given that genetic alterations did not explain effects of *RB1* loss, the role of transcriptional adaptations was explored. Both *RB1* deficient and control H1975 cells surviving initial OSM exposure as DTP showed increased expression of a quiescent alveolar gene expression program (**Figure 4A, S4A**). Initial transcriptional responses to OSM were thus similar in *RB1* deficient and control cells. Alveolar gene expression was also induced by erlotinib treatment of PC9 cells indicating the transcriptional response was not cell line or drug specific (**Figure S4B**)^30^. Induction of this quiescent alveolar gene expression program has been proposed to promote survival of lung cancer cells during EGFR TKI treatment^31,33,42^. Over time on OSM, alveolar gene expression declined and mesenchymal gene expression increased, with mesenchymal gene expression increasing earlier and to a greater extent in *RB1* deficient cells (**Figure 4A-B, S4A-F**). Gene set enrichment analysis indicated that the Hallmark EMT gene set was the most enriched among genes differentially expressed between OSM DTEP and TN cells (**Figure S4D-E**). LUAD patient specimens exhibited a similar pattern, alveolar gene expression increased in residual disease (RD) but then declined in progressive disease (PD) coincident with increasing mesenchymal gene expression (**Figure 4C)**^33^. To test if this alveolar epithelial to mesenchymal transition (EMT) was required for OSM resistance, cells were treated with vitamin D (vitD), a known lung epithelial differentiation agent that can suppress EMT ^43–47^. As expected, H1975 cells were responsive to vitD and vitD treatment suppressed EMT in OSM DTEP cells (**Figure S4G-I**). In *RB1* deficient cells, vitD treatment re-sensitized DTEP cells to OSM, suppressed accelerated OSM DTEP progression, and inhibited OSM DTEP colony formation (**Figure 4D-F**). Thus acquired OSM resistance in these LUAD cells is functionally dependent on dynamic and reversible transcriptional changes suppressed by vitD.

**Figure 4:**
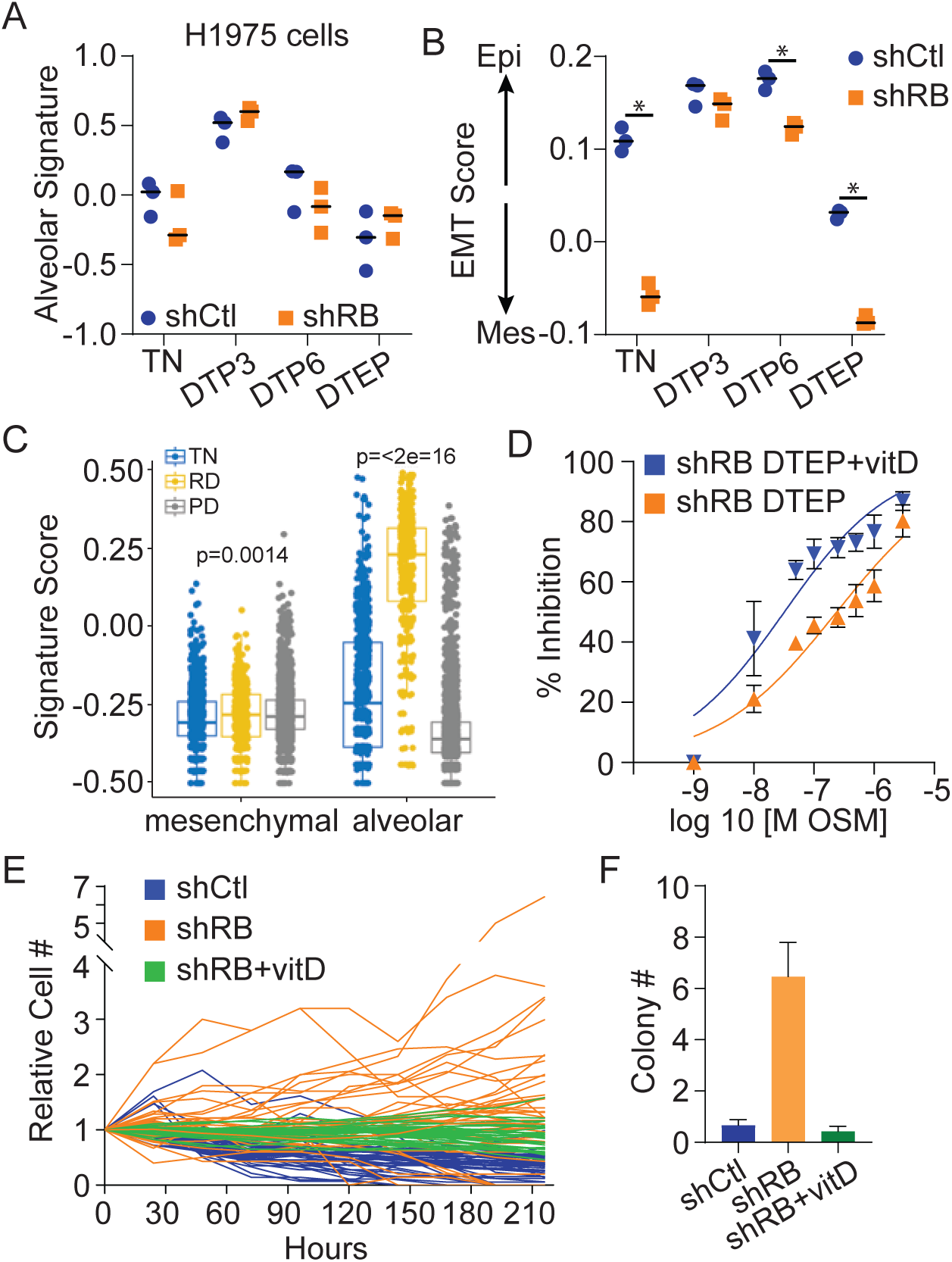
Reversible changes in lineage specific gene expression drive acquired OSM resistance. A) RNA was extracted from the indicated H1975 cells prior to treatment (TN), after 3 days of OSM treatment (DTP3), after 6 days of OSM treatment (DTP6), or at DTEP and analyzed by RNA-seq. A lung alveolar gene expression score was calculated for each of three replicates per sample and plotted with the bar representing the mean. B) An EMT gene expression score was calculated for each sample in A). Asterisks indicate significant differences between shCtl and shRB samples (T-test P<0.05). C) Mesenchymal and lung alveolar gene expression signature scores were calculated from single cell RNA-seq data obtained from lung cancer patient specimens collected prior to treatment (TN), at residual disease during treatment (RD), or at progressive disease during treatment (PD). Data originally generated in Maynard et al. D) RB1 deficient H1975 OSM DTEP cells treated with vitamin D or vehicle were tested for OSM sensitivity in dose response experiments. The percent OSM mediated growth inhibition is plotted versus OSM concentration and non-linear least squares fit of the data plotted. E) The indicated H1975 cells were plated in multi-well plates and treated with OSM or vitamin D+OSM and effects on the outgrowth of resistant cells in each well determined by video microscopy as in figure 2. F) The indicated H1975 cells were treated with OSM or OSM+vitamin D and effects on OSM resistant colony formation measured as in figure 2.

Interestingly, increased mesenchymal gene expression was detected in *RB1* deficient cells prior to OSM exposure (**Figure 5A, S5A**). Consistent with this, data from clinical samples indicated that mesenchymal gene expression was higher in lung cancers with genetic disruption of the *RB1* pathway (**Figure 5B**)^31^. To assess effects of *RB1* loss on transcriptional heterogeneity, the transcriptomes of >223,000 expanded single cell cloned *RB1* deficient and control H1975 cells were analyzed by scRNA-seq before and during treatment with OSM or vitD (**Figure 5C**). OSM DTEP cells treated with OSM, both *RB1* deficient and control, exhibited gene expression consistent with actively proliferating cells as expected (**Figure S5B,C**). As in bulk RNA-seq data, *RB1* deficient TN cells exhibited increased mesenchymal gene expression relative to control TN cells (**Figure S5D**). Fifteen distinct transcriptional clusters were resolved among the cells analyzed (**Figure 5D**). *RB1* deficient TN cells mapped primarily to four different transcriptional clusters (clusters 2, 5-7) while TN control cells from each of two different clones mapped mostly to a single cluster each (cluster 3 or 4)(**Figure 5E**). scRNA-seq data was analyzed by CytoTRACE2, an algorithm designed to quantitate transcriptional plasticity and developmental potential^48^. After filtering for G1 cells to control for cell cycle effects, mean CytoTRACE2 scores were higher in *RB1* deficient TN cells than in control TN cells with *RB1* loss generating a larger fraction of cells with high plasticity (**Figure 5F**). CytoTRACE2 scores declined as *RB1* deficient cells progressed to OSM resistant DTEP, consistent with transcriptional cluster 10 becoming dominant in these DTEP cells (**Figure 5E-F**). Similar results were observed in clinical specimens. Cells with gene expression indicative of *RB1* loss of function exhibited higher CytoTRACE2 scores (**Figure 5G**). CytoTRACE2 scores also increased in clinical specimens as disease progressed through EGFR TKI (**Figure 5H**). Most of the top differentially up regulated genes in OSM resistant cluster 10 have been implicated previously in EGFR TKI resistance and/or EMT (**Table S7**). Control DTEP cells mapped almost exclusively to transcriptional cluster 12 that showed up regulation of an overlapping subset of EGFR TKI resistance associated genes (**Figure 5I**).

**Figure 5:**
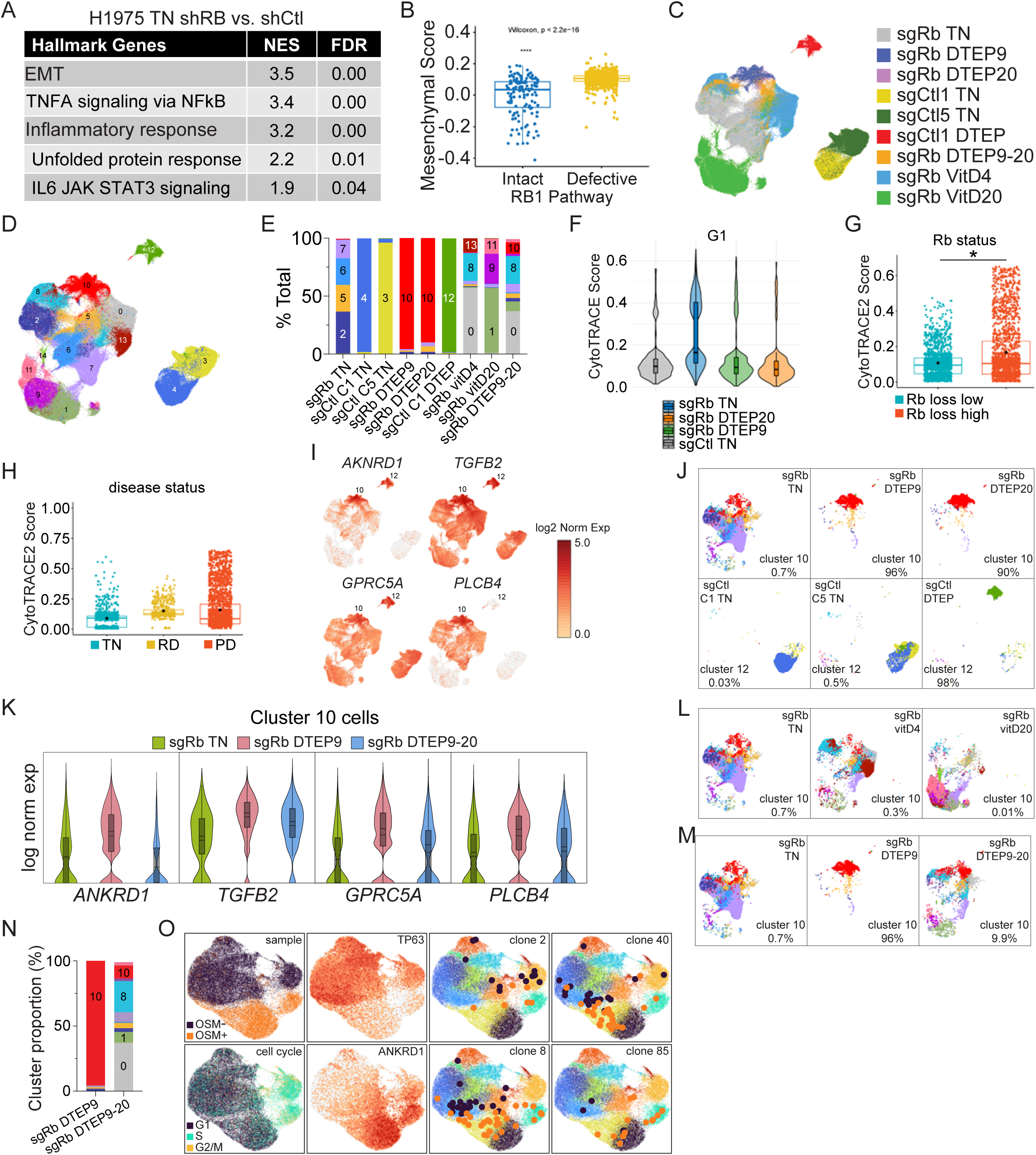
RB1 deficiency drives transcriptional plasticity enabling acquisition of reversible, OSM resistant transcriptional states. A) RNA-seq data from TN *RB1* deficient or control H1975 cells was compared by gene set enrichment analysis using the MsigDB Hallmark gene sets. The top five gene sets enriched in TN *RB1* deficient cells are listed. B) The mesenchymal gene expression score was calculated from scRNA-seq data from patient specimens segregated by the genomic status of the RB1 pathway (*RB1, CDKN2A, CCND1-3, CDK4,6*), indicating mesenchymal gene expression was significantly higher in specimens with RB1 deficiency. Data originally generated in Maynard et al. C) Cells with the indicated genotypes and treatment were analyzed by scRNA-seq, transcriptionally clustered, and UMAP projected with cells color coded by sample identity. D) UMAP projected scRNA-seq data in C) color coded by distinguishable transcriptional clusters. E) The composition of each listed sample based on identified transcriptional clusters from D) are plotted as percent of total cells. F) scRNA-seq data from the indicated samples was analyzed by CytoTrace2 with the distribution of CytoTrace2 scores for each cell displayed in violin plots. G) CytoTRACE2 was used to calculate scores from the clinical scRNA-seq data from Maynard et al., with cells segregated by an RB1 loss gene expression signature score either above (high=low RB1 activity) or below (low=high RB1 activity) the median score. The scores are displayed in box plots. H) Clinical data was analyzed by CytoTRACE2 as in G) with cells segregated by disease stage (treatment naïve=TN, residual disease=RD, progressive disease=PD). I) The UMAP in C) is color coded based on log2 normalized expression of the indicated genes. OSM resistance associated clusters 10 and 12 are highlighted. J) The UMAPs for the indicated samples are displayed color coded by the transcriptional clusters as in D). The fraction of total cells within resistance associated transcriptional clusters are noted. K) The distribution of log normalized expression of the indicated genes among cluster 10 cells from the listed samples is displayed in violin plots. L) UMAPs for sgRb TN cells, sgRb TN cells treated with vitD for 4 days (sgRb vitD4), or sgRb TN cells treated with vitD for 20 days (sgRb vitD20) are shown color coded by transcriptional clusters as in D). The fraction of cluster 10 cells within each sample is shown. M) UMAPs for the indicated samples are shown color coded for transcriptional clusters in D). DTEP9-20 represents RB1 deficient cells treated for OSM for nine days, then OSM is removed for the subsequent 11 days before analysis. The fraction of cluster 10 cells within each sample is indicated. N) The relative proportion of cells mapping to transcriptional clusters in D) is shown for the indicated samples. O) A single cell clone of sgRb H1975 cells was expanded, treated with OSM (1 µm) for 9 days, then either maintained on OSM (OSM+) or OSM removed for the next 11 days (OSM-). Cells were analyzed by scRNA-seq and data displayed in UMAPs color coded by sample, cell cycle phase, expression of OSM resistance (*ANKRD1*) or sensitivity (*TP63*) associated markers, or transcriptional clusters as in D). Each of the four UMAPs at right identify cells within a single clonal lineage with larger dots color coded by sample identity.

Importantly, a small fraction of TN *RB1* deficient cells (0.7%) map to transcriptional cluster 10 prior to OSM exposure (**Figure 5J**). Clone C1 control cells had a much smaller fraction of TN cells mapping to their respective resistance associated transcriptional cluster 12 (0.03%), concurrent with their longer time to OSM DTEP progression (**Figure 1D**). In contrast, clone C5 TN control cells with accelerated time to OSM progression comparable to *RB1* deficient clones (**Figure S1H, S2C**) exhibited an increased fraction of cluster 12 cells (0.5%). *RB1* deficient TN cluster 10 cells express the same EGFR TKI resistance associated genes, but at lower levels compared to their OSM selected DTEP counterparts (**Figure 5K**). Thus rare *RB1* deficient cells spontaneously assume a transcriptional state (cluster 10) that is primed for OSM resistance. The time lag between OSM exposure and resumption of cell proliferation (**Figure 2**) and the difference in frequency between cluster 10 cells (0.7%) and DTEP colony forming cells (0.02%) indicated primed cells don’t become fully OSM resistant until exposure to OSM. The rate of acquired OSM resistance, therefore, correlated with the fraction of cells exhibiting resistance primed transcriptional states prior to treatment.

*RB1* deficient TN cells were treated with vitD to test its impact on resistance associated cluster 10 cells. VitD treatment reduced the fraction of TN cluster 10 cells from 0.7% to 0.01% in a time dependent fashion (**Figure 5L**), coincident with the ability of vitD to suppress accelerated progression through OSM (**Figure 4**). An eleven-day drug holiday (DTEP9-20) also reduced the proportion of OSM resistance associated cluster 10 cells and reduced expression of OSM resistance associated genes within those cells (**Figure 5E, M-N**). This drug holiday induced reduction in cluster 10 cells re-sensitized DTEP cells to OSM in dose response experiments (**Figure S5E-F**). The reversible cluster 10 transcriptional state was thus required for acquired OSM resistance.

Lineage plasticity within individual cells could account for observed transcriptional changes during OSM exposure and drug holidays, but they could also be due differential expansion of genetic subclones in the presence or absence of OSM. To distinguish between these possibilities, *RB1* deficient cells were exposed to OSM for nine days to select for DTEP, tagged with expressed lineage barcodes^49^, and OSM was then either maintained or removed for the next 11 days. Single cell transcriptomes were analyzed to identify clonal cell lineages and trace their transcriptional states. *RB1* deficient DTEP cells reproduced the increased expression of OSM resistance associated genes (e.g. *ANKRD1*) as expected, and transcriptional states shifted during the drug holiday to those characteristic of OSM sensitivity (e.g. *TP63*)(**Figure 5O**). Cells within the same clonal lineage assumed multiple OSM resistant states in the presence of OSM, but shifted to multiple OSM sensitive states during the drug holiday. CytoTRACE2 analysis demonstrated that transcriptional diversity within clonal lineages was not significantly different from the cell population as a whole (**Figure S5G**). Thus lineage plasticity within individual cells, rather than differential expansion of genetic subclones, accounts for the transcriptional adaptations observed. Overall, these observations indicated that *RB1* deficiency drives increased transcriptional plasticity within *EGFR* mutant LUAD cells, increasing the probability rare cells spontaneously assume reversible, OSM resistance primed states that rapidly progress to DTEP during OSM exposure.

### Tumor suppressor gene deficiency drives LUAD lineage plasticity and EGFR independence in vivo

Doxocycline (Dox) inducible expression of the clinically relevant L858R *EGFR* mutant in alveolar type 2 (AT2) cells of the mouse causes LUAD^50^. Suppression of L858R EGFR expression upon Dox withdrawal results in complete and sustained tumor regressions, demonstrating resulting tumors remain dependent on mutant *EGFR* signaling. We have adapted this mouse model to study effects of tumor suppressor gene (TSG) loss in vivo. Two approaches are used to link L858R EGFR expression with TSG deletion. Tet-EGFR-2F mice include a TetO-Cre transgene^51^ and floxed *Rb1/Trp53* alleles^52^ so that AT2 cell specific expression of the reverse tetracycline transactivator (rtTA) induces both L858R EGFR and Cre expression upon Dox administration (**Figure 6A**). Mice lacking floxed alleles or the TetO-Cre transgene (Tet-EGFR) serve as *Rb1/Trp53* wild type controls. Ad-EGFR-2F or Ad-EGFR-3F mice contain floxed alleles of *Rb1/Trp53* (2F) or *Rb1/Trp53/Pten* (3F)^53^, a transgene expressing rtTA under the control of a lox-stop-lox cassette^54^, and the TetO-L858R EGFR transgene (**Figure 6B**). Administration of adenovirus expressing Cre specifically in alveolar type 2 cells (AT2)^55^ induces expression of rtTA and deletes the TSGs while Dox administration induces L858R *EGFR* expression in TSG deleted cells. These three tumor suppressor genes were deleted in combination due to their known roles in EGFR TKI resistance and histological transformation^12,55–59^.

**Figure 6:**
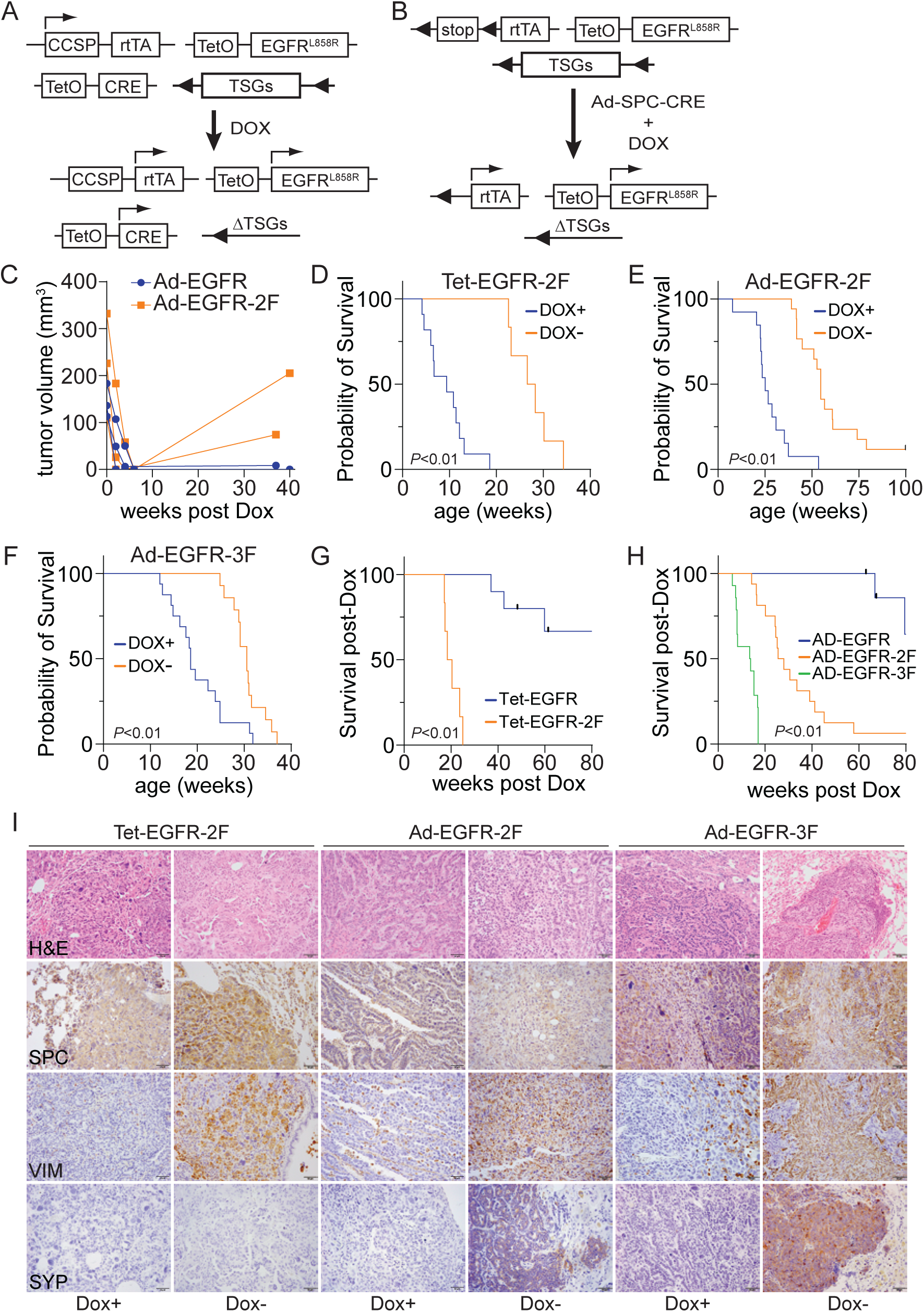
Tumor suppressor gene loss drives progression to mutant *EGFR* independent tumors. A) The schematic shows the design of the Tet-EGFR series of mice that genetically couple doxycycline (Dox) inducible mutant *EGFR* expression with loss of floxed (TSG) in alveolar cells throughout the lung. B) The design shows the Ad-EGFR series of mice that use adenovirus expressed Cre (Ad-SPC-Cre) to trigger Dox inducible mutant EGFR expression and TSG loss focally in lung alveolar cells. C) The response of Ad-SPC-Cre triggered EGFR mutant lung tumors, retaining (Ad-EGFR, N=3) or lacking Rb1/Trp53 (Ad-EGFR-2F, N=3), to removal of Dox as measured by MRI. Total tumor volumes are shown over time with each line representing an individual mouse. D) Overall survival of Tet-EGFR-2F mice in the presence or after removal of Dox once signs of tumor induced morbidity develop. Data is plotted as Kaplan-Meier survival curves that are significantly different by log rank test. E) Overall survival of Ad-EGFR-2F mice lacking Rb1/Trp53 in the presence or absence Dox as in D). F) Overall survival of Ad-EGFR-3F mice lacking Rb1/Trp53/Pten in the presence of absence of Dox as in D). G) Survival of Tet-EGFR (N=6) and Tet-EGFR-2F (N=6) mice post-Dox removal with data plotted as Kaplan-Meier curves as in D). H) Survival of Ad-EGFR (N=9), Ad-EGFR-2F (N=20), and Ad-EGFR-3F (N=14) mice post-Dox removal as in G). I) Representative images of lung tissue sections from mice of the indicated genotype maintained on Dox or relapsing post-Dox removal, stained with H&E or immunostained for the indicated lineage specific marker proteins. Scale bars represent 50 µM.

Pathological evaluation (S.P.) indicates lung tumors arising in Tet-EGFR-2F and Ad-EGFR-2F mice administered Dox are diffuse or nodular adenocarcinomas and giant cell carcinomas exhibiting poor differentiation with solid and papillary growth patterns (**Figure S6A**). Occasional lymphovascular invasion is observed. Liver metastasis is not detected. Control Tet-EGFR mice retaining wild type *Rb1/Trp53* develop nodular and diffuse lung adenocarcinomas showing moderate differentiation, also with solid and papillary growth patterns. Lung tumors developing in all mice express L858R EGFR and markers characteristic of lung adenocarcinoma (**Figure S6A**). Median survival of Tet-EGFR-2F mice is 9.4 weeks from the onset of continuous DOX exposure, less than the 25.3-week median survival of Tet-EGFR mice retaining wild type *Rb1/Trp53* (**Figure S6B**). Median survival of Ad-EGFR-2F (25.1 weeks) and Ad-EGFR-3F (18.3 weeks) mice is also shorter than Ad-EGFP control mice retaining wild type tumor suppressor genes. Tumor suppressor gene loss thus accelerates L858R EGFR induced primary lung tumor progression, consistent with prior findings^60^.

Once mice exhibited early signs of lung cancer morbidity, Dox was withdrawn to suppress L858R EGFR expression and model therapy by testing the tumors’ dependence on continued mutant EGFR activity. Dox withdrawal was used rather than EGFR TKI treatment because EGFR TKI resistance in these mice is often caused by drug specific second site *EGFR* mutations^61,62^ that would preclude assessment of off-target resistance mechanisms. Dox withdrawal elicited complete tumor responses by 4-6 weeks post-Dox in both Ad-EGFR-2F and Ad-EGFR control mice as measured by MRI (**Figure 6C**), so TSG loss did not significantly impact initial responses to therapy. Tumor re-growth was detectable in Ad-EGFR-2F mice, but not in control Ad-EGFR mice, by 40 weeks post-Dox withdrawal. Progression to lethal disease occurred in most Ad-EGFR-2F, Ad-EGFR-3F, and Tet-EGFR-2F mice in the absence of Dox, although they lived significantly longer than mice maintained on Dox (**Figure 6D-F**). The median survival on therapy was 28 weeks for Ad-EGFR-2F mice, 13.7 weeks for Ad-EGFR-3F mice, and 19.4 weeks for Tet-EGFR-2F mice (**Figure 6G-H**). In contrast, tumors developing in Tet-EGFR and Ad-EGFR control mice did not reach median survival by 80 weeks post-Dox. Lung tumors were not detected histologically in lung tissue from a control mouse examined at 20 weeks post-Dox consistent with a complete response. A lung tumor was detected in a control mouse examined at 43 weeks post-Dox (**Figure S6C**) consistent with disease progression to lethal disease in the absence of DOX given sufficient time.

End stage tumors relapsing in the absence of Dox were histologically and molecularly heterogeneous. As expected, L858R EGFR expression was low or undetectable in tumors relapsing post-Dox withdrawal, relative to tumors maintained on Dox (**Figure S6D**). Relapsing tumors retained detectable expression of AT2 cell markers (SPC+) consistent with the cell of cancer origin, but some relapsing tumor foci also expressed proteins characteristic of alternative lineage states like mesenchymal (VIM+) or neuroendocrine (SYP+)(**Figure 6I**). The penetrance of relapsed neuroendocrine tumors was lower in Ad-EGFR-2F mice (1 of 7) than in Ad-EGFR-3F mice (12 of 12), consistent with prior data^56,58^. In contrast the penetrance of relapsed tumors expressing mesenchymal markers was higher in Ad-EGFR-2F mice (4 of 7) than Ad-EGFR-3F mice (5 of 12). Mice maintained on Dox had a lower penetrance of lineage variant tumors (Ad-EGFR-3F 4 of 13 SYP+, 1 of 13 VIM+; Ad-EGFR-2F 0 of 10 NE+, 1 of 10 VIM+)(**Figure 6I**). Lineage variant tumor cells (e.g. VIM+) were more proliferative in the absence of DOX than neighboring non-variant tumor cells (**Figure S6E**), indicating they were less dependent on continued mutant EGFR signaling for proliferation.

Lung cancer tissue sections were profiled by spatial transcriptomics to assess lineage plasticity in an unbiased manner. Integrated analysis of lung tissue sections from four mice identified 12 distinct transcriptional clusters (**Figure 7A-B, S7**). Histologically non-neoplastic clusters (clusters 3-5,11) were shared among all samples. Tissue from a Tet-EGFR+Dox control mouse exhibited only one tumor associated transcriptional (cluster 7). Cluster 7 was the most similar transcriptionally to non-neoplastic epithelial cells, consistent with the more differentiated LUAD histology in this genotype. Tissue from the Tet-EGFR-2F+Dox mouse contained 3 additional neoplastic transcriptional clusters. Relapsing Tet-EGFR-2F-Dox neoplastic tissue exhibited the greatest number of distinct transcriptional clusters, at least 8. A reference single cell RNA-seq dataset from normal mouse lung was used to deconvolve spatial data to cell type specific gene expression signatures (**Figure S8A-B**). Most tumor foci exhibited AT2 cell-like gene expression as expected given the engineered cell of cancer origin (**Figure S8C**). However, relapsed Tet-EGFR-2F-Dox tumor foci also exhibited gene expression characteristic of other lung cell types including club or AT1 cells.

**Figure 7:**
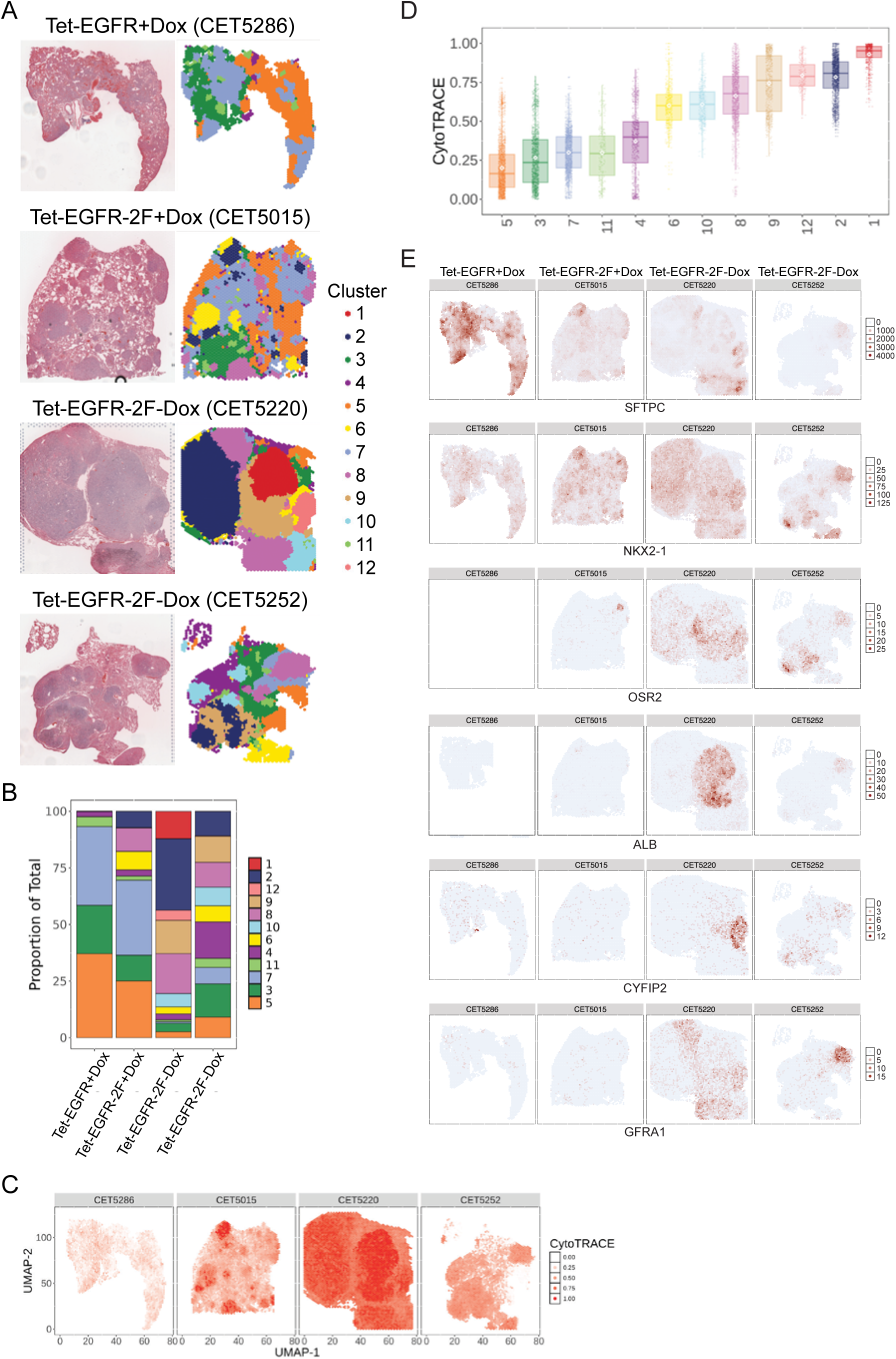
Loss of tumor suppressor genes drives lineage plasticity during L858R EGFR suppression in vivo. A) Lung tissue sections from mice of the indicated genotypes and treatment were stained with H&E (images at left) and evaluated by spatial transcriptomic profiling. Integrated analysis of the four sections identified 12 distinguishable transcriptional clusters (images at right color coded by transcriptional cluster). B) The composition of each sample based on the relative proportion of each detected transcriptional cluster is displayed highlighting that *Rb1/Trp53* deficient tumors progressing through Dox withdrawal exhibit increased transcriptional complexity. C) CytoTRACE2 scores are calculated for each transcriptionally profiled spot and displayed in a color code proportional to the score. CytoTRACE2 scores increase upon *Rb1/Trp53* loss and suppression of mutant EGFR expression. D) The distribution of spot CytoTRACE2 scores is plotted for each transcriptional cluster as box plots. E) Sections are shown with transcriptional profiled spots color coded based on the relative expression of the indicated lineage specific genes.

CytoTRACE2, was used to quantify transcriptional plasticity^48^. Neoplastic transcriptional clusters exhibited higher CytoTRACE2 scores compared to non-neoplastic transcriptional clusters (**Figure 8C-D**). Tumor foci arising in Tet-EGFR+Dox control mice exhibited the lowest neoplastic CytoTRACE2 score (cluster 7) and showed the highest expression of the AT2 cell specific gene *Sftpc* (**Figure 7E**), consistent with their differentiated histology. The highest CytoTRACE2 scores were observed in relapsing tumor foci from Tet-EGFR-2F-Dox mice, concordant with the increased number of distinguishable transcriptional clusters detected (**Figure 7D-E**). Some of tumor foci relapsing in the absence of Dox expressed genes normally restricted to cell lineages from other tissues like *Alb* (liver), *Gfra1* (brain), *Cyfip2* (nervous system), or *Osr2* (mesenchymal tissue). Such lineage plasticity was low or absent in Tet-EGFR-2F mice maintained on Dox. Thus tumor suppressor gene deficient LUAD treated to suppress mutant EGFR activity had expanded transcriptional and lineage plasticity compared to control LUAD retaining these tumor suppressor genes or to LUAD retaining mutant EGFR signaling. While prior results have established a role for *Rb1/Trp53* loss in SCLC transformation of EGFR mutant LUAD^24^, results here indicate lineage plasticity facilitated by loss of these tumor suppressor gene extends beyond SCLC to other lineage states with reduced dependence on EGFR signaling. The incidence of treatment associated SCLC, therefore, likely underestimates the impact of lineage plasticity on acquired EGFR TKI resistance.

## Discussion

By leveraging in vitro, in vivo, and clinical data, this study demonstrates that transcriptional plasticity and ensuing transcriptional adaptations can drive acquired EGFR TKI resistance in LUAD. Lineage tracing and scRNA-seq analysis demonstrates these resistance-conferring transcriptional adaptations are dynamic, meta-stable, and reversible. The transcriptionally plastic LUAD state is associated with increased EMT and inflammatory gene expression, but only a small fraction of these plastic cells become EGFR TKI resistant once they express sufficient levels of EGFR TKI resistance conferring genes. The lung epithelial differentiating agent Vitamin D suppresses EMT overall, reduces the fraction of LUAD cells specifically within the EGFR TKI resistant transcriptional state, and inhibits the rate of acquired EGFR TKI resistance. EMT and inflammatory gene expression is thus required, but not sufficient, for EGFR TKI resistance. Consistent with findings in prostate cancer^63,64^, this EMT/inflammatory state may represent a necessary intermediate state from which EGFR TKI resistant cells arise.

The current paradigm for treating acquired EGFR TKI resistance in LUAD is focused on targeting genetic resistance mechanisms, but genetic resistance mechanisms have been identified in only about half of the cases examined. Findings reported here suggest there is an opportunity to improve EGFR TKI outcomes by targeting such non-genetic adaptations^25,65,66^. Therapeutic strategies aimed at stabilizing or inducing the EGFR TKI sensitive epithelial lineage state, like vitD supplementation reported here, may delay progression on EGFR TKI therapy. VitD supplementation has failed in prior clinical trials of unselected LUAD patients, but recent evidence suggests beneficial effects of vitD are confined to *EGFR* mutant LUAD and not KRAS mutant LUAD^36,46,67^. Targeting FAK-YAP signaling that promotes the quiescent alveolar lineage adaptation that allows some LUAD cells to survive therapy initially as DTP^31^ using available YAP^68^ or FAK^69^ inhibitors is another possible approach. The identification of transcriptional programs or genetic alterations like *RB1* loss that are associated associated with transcriptional plasticity provide a framework for developing biomarkers to better predict EGFR TKI responses and guide personalized treatment strategies.

While Rb is a well-established regulator of the cell cycle and genome stability, we demonstrate here that Rb loss accelerates progression through EGFR TKI by increasing transcriptional plasticity within LUAD cells. This plasticity enables treatment naïve LUAD cells to access a greater range of transcriptional states, with rare cells assuming a state primed for EGFR TKI resistance. This primed state is characterized by increased expression of multiple genes that have been functionally linked to EGFR TKI resistance. Upon exposure to therapy, a fraction of primed cells rapidly induce sufficient expression of these genes to proliferate in the presence of clinically relevant EGFR TKI concentrations. Rb loss increases the number of these primed cells thereby accelerating progression of LUAD through therapy. *RB1*-loss thus represents a unique genetic alteration that does not confer EGFR TKI resistance directly but enables therapeutic resistance indirectly by enabling rapid epigenetic adaptations. Epigenetic adaptations may allow LUAD cells to proliferate sufficiently during therapy to either seed disease progression directly or to facilitate de novo acquisition of resistance conferring genetic mutations in LUAD where they do not pre-exist. We suggest this explains, at least in part, why LUAD with loss of function *RB1* mutations exhibit inferior outcomes on EGFR TKI therapy^12^ and why CDK4/6 inhibitors (CDK4/6i) that activate protein can improve EGFR TKI responses^70^. Transcriptional plasticity driven by *RB1* loss is also affect other aspects of cancer progression that rely on adaptation to selective pressue, like metastasis^71,72^.

Questions remain regarding the molecular mechanisms underlying LUAD transcriptional plasticity, but some clues can be ascertained from RB1. RB1 protein (Rb) is well known to physically interact with both transcription factors and chromatin regulatory complexes to mediate epigenetic control of gene expression, and loss of this control could drive transcriptional plasticity. Further study is required to characterize chromatin changes induced by Rb loss and to identify the transcription factors and chromatin modifying complexes mediating effects of Rb loss in the context of *EGFR* mutant LUAD. Elucidating relevant mechanisms could uncover new opportunities for therapeutic intervention to limit transcriptional plasticity and acquired EGFR TKI resistance.

This study highlights the less appreciated role of non-genetic adaptations in accelerating acquired resistance to EGFR TKI. It also suggests that some genetic alterations, like disruption of *RB1*, may facilitate acquired EGFR TKI resistance indirectly by driving transcriptional plasticity. Increased focus on non-genetic molecular mechanisms contributing to the dynamic process of acquired therapeutic resistance is likely to open new avenues for therapeutic intervention, particularly for patients progressing on EGFR TKI for which genetic mechanisms of resistance have not been identified.

## Methods

### Cell culture

EGFR-mutant lung adenocarcinoma cell lines H1975 and HCC827 were obtained from ATCC. The PC9 cell line was generously provided by S.I. Abrams, PhD (RPCCC). H1975 and HCC827 cells were cultured in RPMI 1640 supplemented with 10% fetal bovine serum (FBS) and 1% penicillin-streptomycin. PC9 cells were cultured in RPMI 1640 supplemented with 10% FBS, 1% penicillin-streptomycin (P/S), 7.5% HEPES, 1% L-glutamine, 1% sodium pyruvate, and 1% MEM non-essential amino-acids. Osimertinib or Erlotinib resistant cell lines were generated by culturing cells in growth media containing either 1μM Osimertinib (S7297, selleckchem) or 1μM Erlotinib (CT-EL002, chemietek) for over 40 days. Resistant cells were maintained in Osimertinib/Erlotinib containing media. All cells were cultured at 37°C and 5% CO2. Drugs were dissolved in DMSO as recommended by the manufacturer.

### Lentivirus infection

All lentivirus were produced by Gene Modulation Shared Resource at Roswell Park Comprehensive Cancer Center. Lentiviral plasmids were co-transfected with psPAX2 (*gag, pol, env*) and pMD2.G (VSV-G envelope) into HEK293T cells using LipoD293 (SignaGen) according to the manufacturer. Virus containing media is collected at 32°C for 48-72 hours after transfection. Cells to be infected were plated in 6-well plates the night before infection. The next day, existing media was removed and 500ul viral supernatant + 500ul normal media + 4ug/ml polybrene were added to each well and incubated for 30 minutes at 37°C/5% CO2. 1ml of additional normal media was added and the plates centrifuged at 1800 rpm for 45 minutes at room temperature and then incubated overnight at 37°C/5% CO2. Media was changed to normal media the next morning. RB1 targeted or control guide RNA were cloned into the pL-CRISPR.EFS.tRFP vector^73^ as described^74^ (generously provided by R. Bremner, U. Toronto).

### Growth rate and dose response assays

3000 cells per well were seeded in 12 well plate the night before and fixed for SRB assay at time 0, 24 and 48 hours later. Results of SRB assays were normalized to day 0 to plot the growth curve. Dose response experiments were performed similarly in 12 well plates. The day following plating media was changed and supplemented with increasing concentrations of drug (e.g. OSM). Cells were treated for 9 days and media was changed every 3 days with SRB assay used to measure viable cells post-treatment. IC50 values were determined by non-linear regression of the resulting dose response data.

### Sulfarhodamine B (SRB) Assay

SRB assays were performed by first fixing cells with 10% trichloroacetic acid (TCA) (Thermo Fisher Scientific), washed and air dried, then stained with 0.57% (wt/vol) SRB dye (Sigma-Aldrich), stained cells were washed and air dried, cells solubilized with 10mM Tris-base solubilization buffer, and the optical density of the supernatant read at 510 nm on a BioTek SYNERGY microplate reader.

### Colony formation

500 or 800 cells were plated per well in 6 well plate and cells treated with vehicle or the indicated concentration of drug for the listed number of days, with media changes every 3 days. Cells were then fixed and stained with crystal violet (0.5% w/v)(Sigma-Aldrich), washed and air dried, then counted manually.

### Time lapse imaging

1000 cells were seeded in each well of a 96 well plate and maintained in culture media supplemented with 1uM Osimertinib with media changes every 3 days. Images of each well were captured every 8 hours over 9 days using an Incucyte S3 live cell imager. Cell number and cell division events per well per image were identified manually.

### Western blot analysis

Cell lysates were prepared using RIPA buffer containing a protease and phosphatase inhibitor cocktail (Roche). Lysates were collected into microfuge tubes and incubated on ice for 30-40min with vertexing every 10min. After centrifugation at max speed for 15min at 4°C, supernatant was transferred into clean tubes and protein concentration measured by Bradford assay. 50 ug total protein was loaded into the wells of the SDS-PAGE gel, then transferred from gel to a nitrocellulose membrane. Membrane was blocked in 5% non-fat milk for 1 h at room temperature and incubated with primary antibody overnight at 4°C, followed by incubation with the secondary antibody for 1 h at room temperature. The signal was visualized using a chemiluminescent substrate kit (BioRad).

### Real time qRT-PCR

RNA was extracted using TRIzol (Thermo Fisher Scientific) following the manufacture’s protocol. cDNA was synthesized from RNA using iScript™ cDNA Synthesis Kit (1708890, BioRad). The iQ SYBR Green Supermix (1708880, BioRad) and CFX Connect system (Bio-rad) were used for qRT-PCR reactions. Relative mRNA levels were determined using the 2-ΔΔCt method.

### DNA sequencing

Targeted sequencing of *EGFR* Exons 19-21 was performed according to Molina-Vila, et al.^75^ Resulting sequences were aligned against the NCBI Reference Sequence NG_007726.3 using the ClustalW2 alignment tool. For whole exome DNA sequencing, individual target capture for each DNA sample was carried out using the SureSelect XT HS2 DNA Target Enrichment System (Agilent Inc). 100ng of DNA was sheared using a Covaris E220 (Covaris Inc.) to a size range of 150bp-200bp followed by end repair, duplex UMI adaptor ligation, and 10 cycles of PCR to complete the dual-index adapters. Unique dual-indexed, molecular-barcoded libraries were purified with AMPureXP beads (Beckman Coulter) and validated for appropriate size (225-275bp) on a Tapestation 4200 DNA1000 screentape (Agilent Inc.). 1000ng of purified library was then hybridized to the SureSelectXT Human All Exon V7 Capture library (Agilent Inc.). The hybridized regions were then bound to streptavidin magnetic beads and washed to remove any non-specific bound products. Eluted libraries underwent a second 10 cycle PCR amplification to generate enough material for sequencing. Final libraries were purified, measured by Tapestation 4200 DNA1000 screentape (250-350bp), and quantitated using KAPA qPCR (KAPA Biosystems). Individual libraries were pooled in equimolar 2nM final concentration. Each pool is denatured and diluted to 200pM with 1% PhiX control library added. The resulting pool is then loaded into the appropriate NovaSeq Reagent cartridge, as determined by the number of sequencing cycles desired, and sequenced on a NovaSeq6000 following the manufacturer’s recommended protocol (Illumina Inc.).

### RNA-Sequencing

Total RNA purification was performed using the miRNeasy mini kit (Qiagen) following manufacturer’s recommendation, quantified using a Qubit Broad Range RNA kit (Thermofisher), and concentration determined using Ribogreen fluorescent binding. RNA quality is evaluated using RNA Nanotape on the 4200 Tapestation (Agilent technologies). Sequencing libraries are prepared from 500ng total RNA using the RNA HyperPrep Kit with RiboErase (HMR) (Roche Sequencing Solutions) following manufacturer’s instructions. Samples are then purified, fragmented, primed for cDNA synthesis, reverse transcribed into first strand cDNA using random primers, the RNA template is removed, and DNA replacement strand synthesized incorporating dUTP to generate ds cDNA. Pure Beads (KAPA BIOSYSTEMS) are used purify the ds cDNA and a single adenine nucleotide is added to the 3’ ends of the blunt cDNA fragments. Multiple indexing adapters containing a single thymine nucleotide are ligated to the ends of the ds cDNA. Adapter ligated libraries are amplified by PCR, purified using Pure Beads, and validated for appropriate size on a 4200 TapeStation D1000 Screentape (Agilent Technologies, Inc.). The DNA libraries are quantitated using KAPA Biosystems qPCR kit, and are pooled in an equimolar fashion, following experimental design criteria. Each pool is denatured and diluted to 350pM with 1% PhiX control library added. The resulting pool is then loaded into 200 cycle NovaSeq Reagent cartridge, for 100 cycle paired end sequencing, and sequenced on a NovaSeq6000 following the manufacturer’s recommended protocol (Illumina Inc.).

### scRNA-seq

Pools of Rb-deficient (sgRb) or Rb-proficient (sgCtl) H1975 cells were single cells cloned by limiting dilution, expanded, treated as described, then analyzed by scRNA-seq. For cell lineage tracing, expanded single cell cloned Rb-deficient cells were cultured with OSM (1 μM) for 9 days, infected with lentivirus expressing CellTags^49^, expanded for 3 days in OSM to generate clone progeny, then split into two groups. One group was grown in the presence of OSM and the other in the absence of OSM until day 20.

Single cell gene expression libraries were generated using Evercode Combinatorial Barcoding Technology from Parse Biosystems. Individual transcriptomes are uniquely labeled by passing fixed cells or nuclei through four rounds of barcoding. In the first round of barcoding, fixed cell samples are distributed into 48 wells, and cDNA is generated with an in-cell reverse transcription reaction using well-specific barcoded primers. Cells from each well are pooled and distributed into 96 wells followed by an in-cell ligation reaction that appends a second well-specific barcode to the cDNA. Pooling, distribution and in-cell ligation are repeated for a third round of barcoding. After three rounds of barcoding, the cells are pooled and split into 8 distinct sublibraries. The cells in these sublibraries are lysed and the barcoded cDNA is isolated. A fourth sublibrary-specific barcode is introduced by PCR to each cDNA molecule. The resulting libraries are evaluated on D1000 screentape using a TapeStation 4200 (Agilent Technologies), and quantitated using Kapa Biosystems qPCR quantitation kit for Illumina. They are then pooled, denatured, and diluted to 200pM with 5% PhiX control library added. The resulting pool is then loaded into the appropriate NovaSeq Reagent cartridge and sequenced on a NovaSeq6000 following the manufacturer’s recommended protocol (Illumina Inc.).

### Spatial transcriptomic analysis

Tissue embedded paraffin blocks were sectioned and trimmed to fit within capture areas on the Visium Spatial slides. Deparaffinization and H&E staining was performed, followed by imaging of each tissue sample. 10x Visium library preparation was performed using Visium Spatial Transcriptomic V1 kit following manufacturers recommendations. Briefly, the RNAs within the tissue were hybridized to the mouse whole transcriptome probe panel and the hybridized probes were captured on the Visium slides. Captured probe products were then extended using the unique molecular identifier (UMI), Spatial Barcode, and partial Read 1 adapter to generate spatially barcoded and ligated probe products. The ligated products were released from the slide and qPCR was used to determine sample index PCR cycles. Final libraries were generated by PCR using sample indexing primers. The resulting libraries were evaluated using D1000 Screen Tape on the TapeStation 4200 (Agilent Technologies) and quantitated using the KAPA Biosystems qPCR quantitation kit for Illumina. They were then pooled, denatured, and diluted to 300 pM (picomolar) with 1% PhiX control library added. The resulting pool was then loaded into the appropriate NovaSeq Reagent cartridge followed by sequencing on a NovaSeq6000 according to the manufacturer’s protocol (Illumina Inc., San Diego, CA, USA). Once sequencing was completed, tissue images taken after H&E staining on the Visium slide were used to align the gene expression from the spatial barcodes unique to each location in the capture area during data analysis.

### Bioinformatics

Bulk RNA-seq data was adjusted for batch-effects using ComBat-Seq. DESeq2 was used to identify statistically significant differentially expressed genes between relevant samples. The DESeq2 log2 fold change statistic was analyzed using Gene Set Enrichment Analysis (GSEA, version3.0_BETA). Pathway Analysis was run against MSigDB^76^. The alveolar module score was calculated using the gene signature from Laughney et al.^77^ To quantify relative epithelial and mesenchymal (EMT) transcriptional states of individual samples or cells, the Singscore (v1.18.0)^78^ R package was employed to analyze transcriptomic data using an established EMT signature^79^.

Raw ParseBio single cell sequencing data was processed with split-pipe version 1.4.2 to generate the filtered UMI count matrix. The Seurat R package was used for data normalization, cell cycle regression, dimensional reduction, batch correction, and cell clustering. Clonal lineages were identified following the methodology of Biddy et al.^49^ For scRNA-seq data from clinical samples, read counts and metadata from Maynard A et al.^33^ were used. Expression data were normalized using a log2 transformation of non-zero gene counts plus one [log2(count + 1)]. To quantify transcriptional plasticity at single-cell resolution, we applied CytoTRACE2^48,80^, a deep learning-based computational tool that infers differentiation potential from scRNA-seq data. A subset of annotated tumor cells was extracted for RB1 signature analysis using a gene signature from Witkiewicz et al.^81^, analyzed with the singscore and GSEABase R packages. Genes were ranked, and signature scores were computed for each tumor cell. Tumor cells were stratified into RBsig High (low *RB1* activity) and RBsig Low (high *RB1* activity) groups based on the median signature score.

and visualized these scores per cell and grouped by either experimental treatment, clonal lineage, or genotype. The algorithm built-in log2-adjusted representation of the input expression data was used for prediction of top ranked expression profiles in order to capture transcriptomic signals. The predicted potency scores provide a continuous measure of developmental potential, ranging from 0 (differentiated) to 1 (totipotent) for each cell. The association between CytoTRACE2 scores with RB1 loss signature scores or disease status (TN, RD or PD) was visualized as boxplots. Unpaired Wilcoxon rank-sum test was applied to test for statistical significance between groups and the Kruskal-Wallis test was used for multiple groups test. Plasticity scores were visualized with violin or boxplots, with horizontal line indicating the median. To test for statistically significant differences in CytoTRACE2 scores between clonal and non-clonal lineages, Wilcoxon permutation test was used accompanied by Cliff’s Delta to evaluate effect size.

### Genetically engineered mice

All mouse work was performed under IACUC approved protocols at Roswell Park Comprehensive Cancer Center. Animals are housed on a 12-h light/dark cycle at 22±2°C. All mice were monitored daily for signs of morbidity. CCSP-rtTA;TetO-EGFRL858R mice were described previously^50^ and generously provided by P. Tran (Johns Hopkins University). B6.Cg-Tg(tetO-cre)1Jaw/J^51^ (#006234) and B6.Cg-Gt(ROSA)26Sortm1(CAG-rtTA3)Slowe/LdowJ^54^ (#029617) mice were purchased from The Jackson Laboratory. Use of floxed Rb1, Trp53, and Pten alleles has been described in our prior work^56,71^. Final genotypes are on a mixed C57BL/6:129/Sv:FVB background. Littermates of relevant genotypes are used as test and control mice when available. Mouse tail DNA was collected for PCR genotyping using primers specified in the originating citations noted above. Tails were boiled in 100 μl of 50mM Sodium chloride for 30min, neutralized by addition of 30μl of Tris-HCl (pH 7.5), centrifuged for 1 minute at max speed in a 1.5 ml Eppendorf tube, and supernatants collected for PCR analysis. DNA amplification products were resolved by agarose gene electrophoresis and visualized by EtBr staining. Mouse tissue was collected and flash frozen in liquid N2 for molecular studies. Alternatively, tissue was fixed in 10% neutral buffered formalin or phosphate-buffered 4% paraformaldehyde, paraffin embedded, sectioned, and analyzed by H&E or immunostaining as described previously^56,71,82^. For histological analysis, lungs were inflated by injecting phosphate-buffered 4% paraformaldehyde through the trachea in order to prevent tissue collapse.

### Histology and immunostaining

5-μm thick tissue sections were used for routine H&E staining. For immunostaining, pre-warmed slides were deparaffinized in xylene solutions followed by rehydration in graded ethanol. Antigen retrieval was accomplished by boiling for 20 minutes in 10 mM sodium citrate solution (pH 6.0). Slides were blocked by treatment with 3% hydrogen peroxide followed by treatment with 2.5% normal goat serum (Vector Laboratories) for 60 minutes at room temperature. Primary antibody incubations occurred overnight in a humidified chamber. Slides were washed and incubated for 60 minutes with secondary antibody conjugated to peroxidase (Vector Laboratories). Staining was developed by treatment with diaminobenzidine (Dako, K3468) at room temperature. Immunostained sections were counterstained with hematoxylin prior to dehydration and mounting.

## Acknowledgements

We acknowledge Dineen Vogler for technical contributions to this work. We thank the laboratory of Dominic Smiraglia for helpful discussions. This work was supported by grants from the NCI (R01CA272771 to B.A.; R50CA221786 to M.C.; R01CA234162 and U24CA274159 to D.W.G.; R21CA282428 to E.V.B.; R50CA283805 to P.S.; U54CA224081, R01CA204302, U01CA272546, R01CA238236 to T.B.). This work was supported by the Roswell Park Comprehensive Cancer Center Alliance Foundation and NCI grant P30CA016056 that supports the Roswell Park shared resources utilized in this work.

---

**Figure S1:**
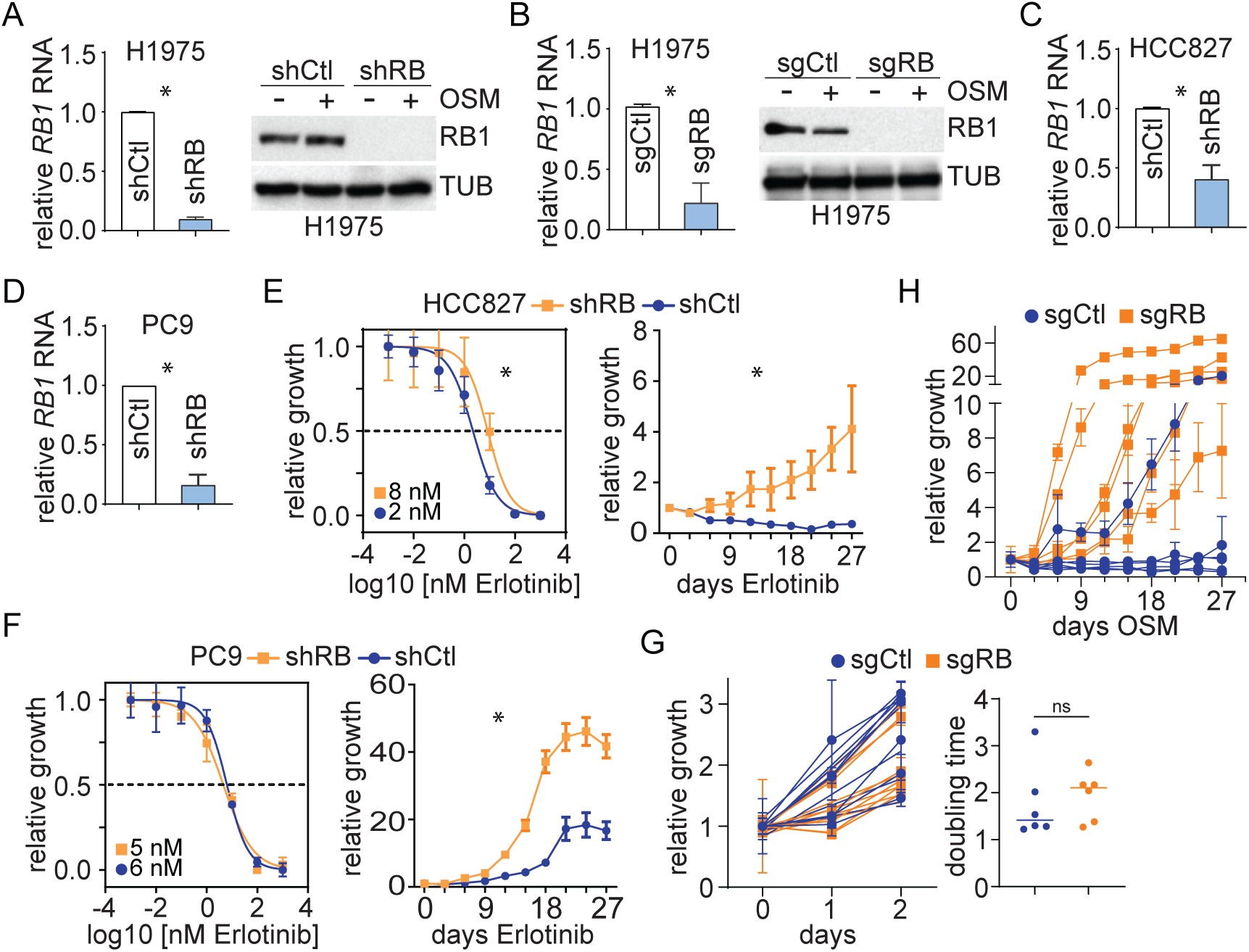
RB1 deficiency accelerates acquired EGFR TKI resistance in human LUAD cell lines. A) RNA and protein were extracted from three replicates of H1975 cells modified by control or *RB1* targeted shRNA, and relative *RB1* RNA and protein levels measured by real time RT-PCR or Western blotting. Asterisks indicate a significant differences between samples (T-test P<0.05). Tubulin serves as the protein loading control. B) H1975 modified by control or *RB1* targeted sgRNA+Cas9 were analyzed as in A). C) RNA was extracted from HCC827 cells modified by control or *RB1* targeted shRNA and relative RB1 RNA analyzed as in A). D) RNA was extracted from PC9 cells modified by control or *RB1* targeted shRNA and relative RB1 RNA measured as in A). E) Erlotinib sensitivity (left panel) and time to resistance (right panel) of the indicated modified HCC827 cells was measured, analyzed, and plotted as in figure 1. F) Erlotinib sensitivity (left panel) and time to resistance (right panel) was measured in modified PC9 cells as in E). G) H1975 cells modified by control or RB1 targeted sgRNA+Cas9 were single cell cloned, expanded, treated with OSM (1 µM), and cell accumulation curves plotted (left panel). Doubling times were calculated from the curves and plotted with their mean value indicated by the line (right panel). Samples are not significantly different by T-test (P>0.05). H) Time to OSM resistance (1 µM) was measured in modified H1975 single cell clones from G). Samples are significantly different by nested 1-way ANOVA (P<0.05).

**Figure S2:**
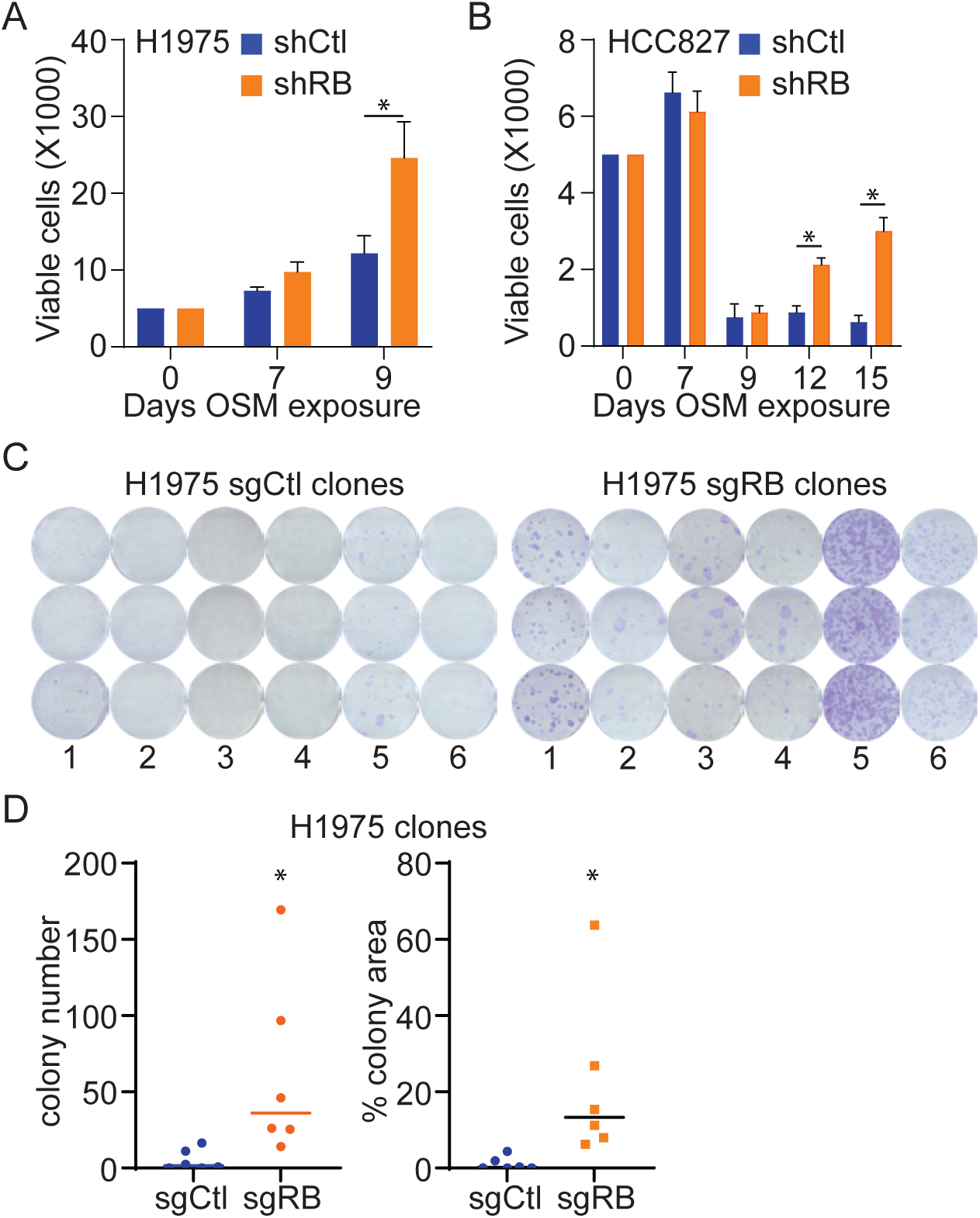
Rare *RB1* deficient LUAD cells rapidly progress to OSM DTEP. A) and B) The indicated cells were plated and exposed continuously to 1 µM OSM. Viable cell accumulation was monitored over time. Asterisks indicate differences between samples are statistically significant (T test P<0.05). C) Single cell clones were derived from the indicated cells, expanded, plated in 6-well plates, and exposed to 1 µM OSM continuously for 24 days to assess OSM resistant colony formation. Representative images from 3 replicates of six clones are shown. D) The mean colony number or colony area from the experiment in C) are plotted for each clone. Asterisks indicated difference between samples are statistically significant (T test P<0.05).

**Figure S3:**
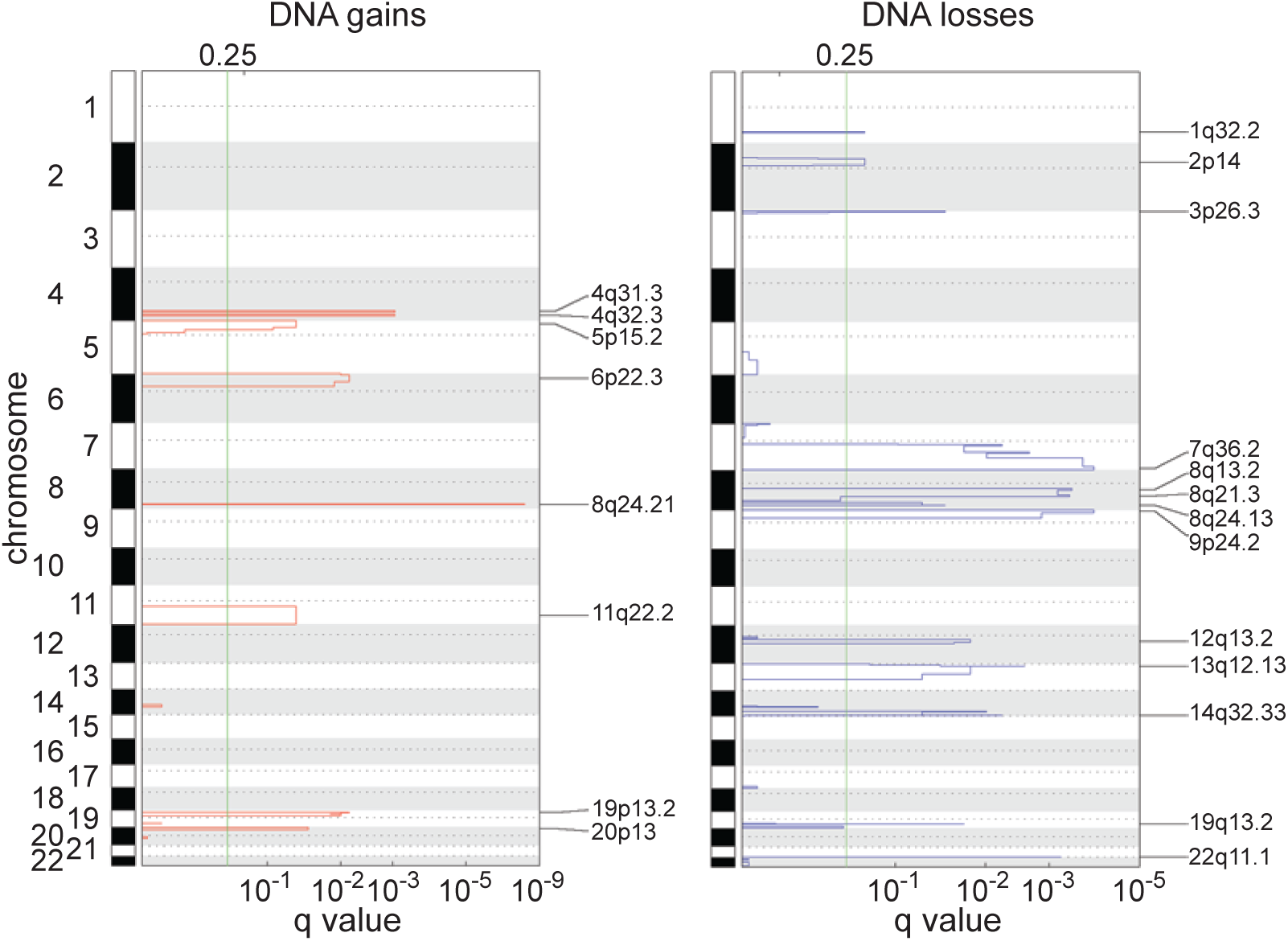
Focal amplification at 8q24.21 spanning the MYC gene detected in OSM DTEP samples. The q value for DNA gains or losses, relative to the H1975 reference genome, for an Rb deficient OSM DTEP sample are shown. The most significant change detected is an amplification at 8q24.21 spanning 16 genes, including *MYC*. This amplification was also detected in RB1 proficient H1975 control OSM DTEP cells.

**Figure S4:**
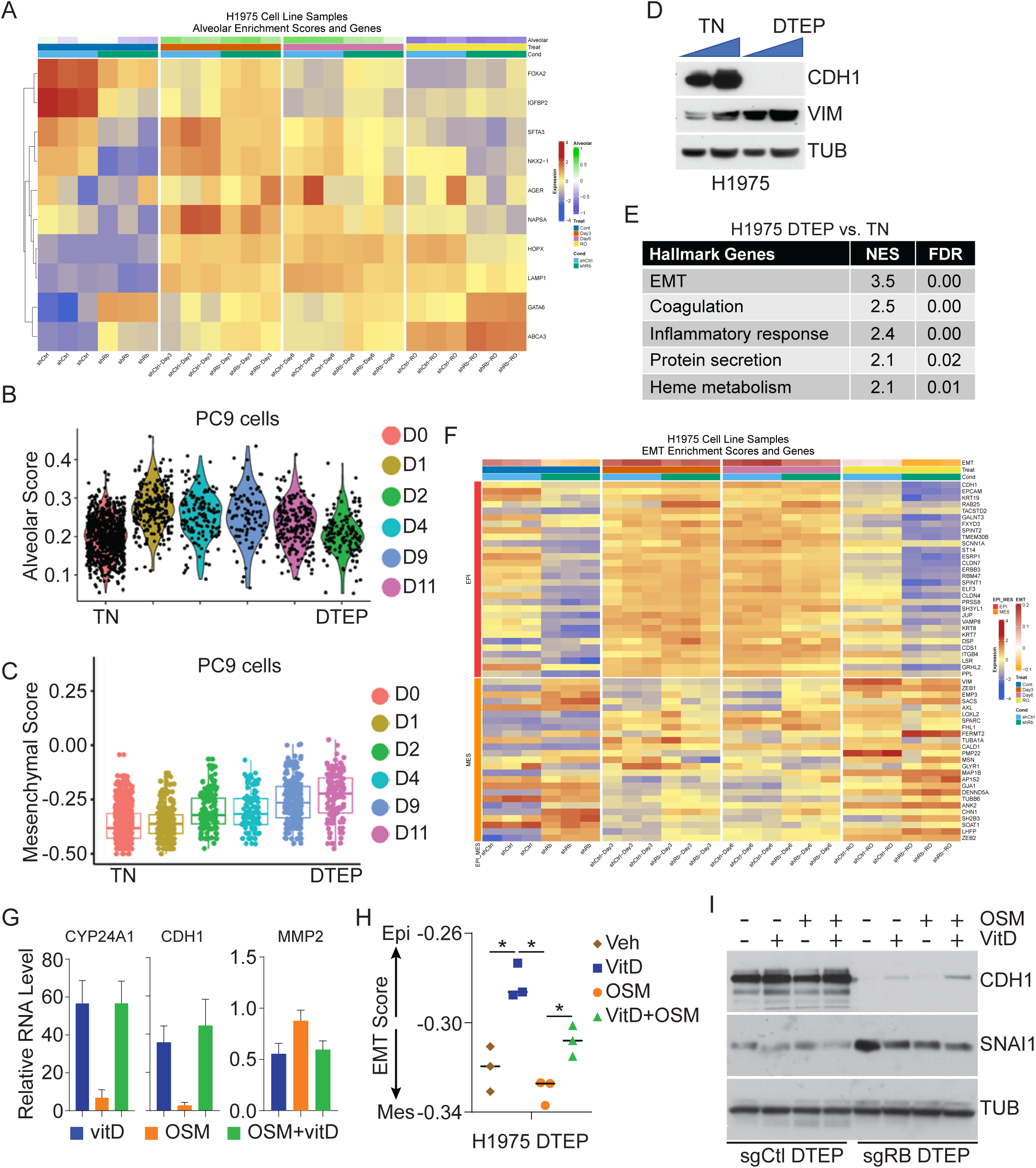
Reversible changes in lineage specific gene expression drive acquired OSM resistance. A) RNA extracted from the indicated H1975 samples was analyzed by RNA-seq and the relative expression of alveolar lineage specific genes shown as a heatmap. The alveolar gene expression signature score is shown at top. B) PC9 cells treated for the indicated number of days with OSM were analyzed by single cell RNA-seq, and the alveolar module scores were calculated for each cell. Data were originally generated in Aissa et al. C) A mesenchymal signature score was calculated from the single cell RNA-seq data in B. D) Protein was extracted from the indicated H1975 cells, and increasing amounts of total protein analyzed for expression of the indicated proteins by western blotting. Tubulin serves as a protein loading control. E) RNA was extracted from the indicated H1975 cells and used to generate RNA-seq data. RNA-seq data was analyzed by gene set enrichment analysis using the MSigDB Cancer Hallmarks gene sets. The five most enriched gene sets differentially expressed in DTEP cells, ranked by NES, are shown. F) Expression of EMT related genes was plotted as a heat map from RNA-seq data in A) with epithelial genes at top and mesenchymal genes at bottom. The EMT gene expression signature score is shown at top. G) H1975 DTEP cells were treated with OSM and/or vitD as indicated (N=3), RNA extracted, and expression of the indicated vitD responsive, epithelial, and mesenchymal marker genes measured by quantitative RT-PCR. The mean and standard deviation of relative expression is plotted. H) RNA from samples in G) were analyzed by RNA-seq and the EMT score calculated. Each dot represents a biological replicate with the bar signifying the mean. The asterisks indicate statistically significant differences between the indicated samples (T-test P<0.05). I) Protein was extracted from the listed H1975 cells treated as indicated, and the indicated epithelial (CDH1) and mesenchymal (SNAI1) marker proteins measured by western blotting. Tubulin serves as the protein loading control.

**Figure S5:**
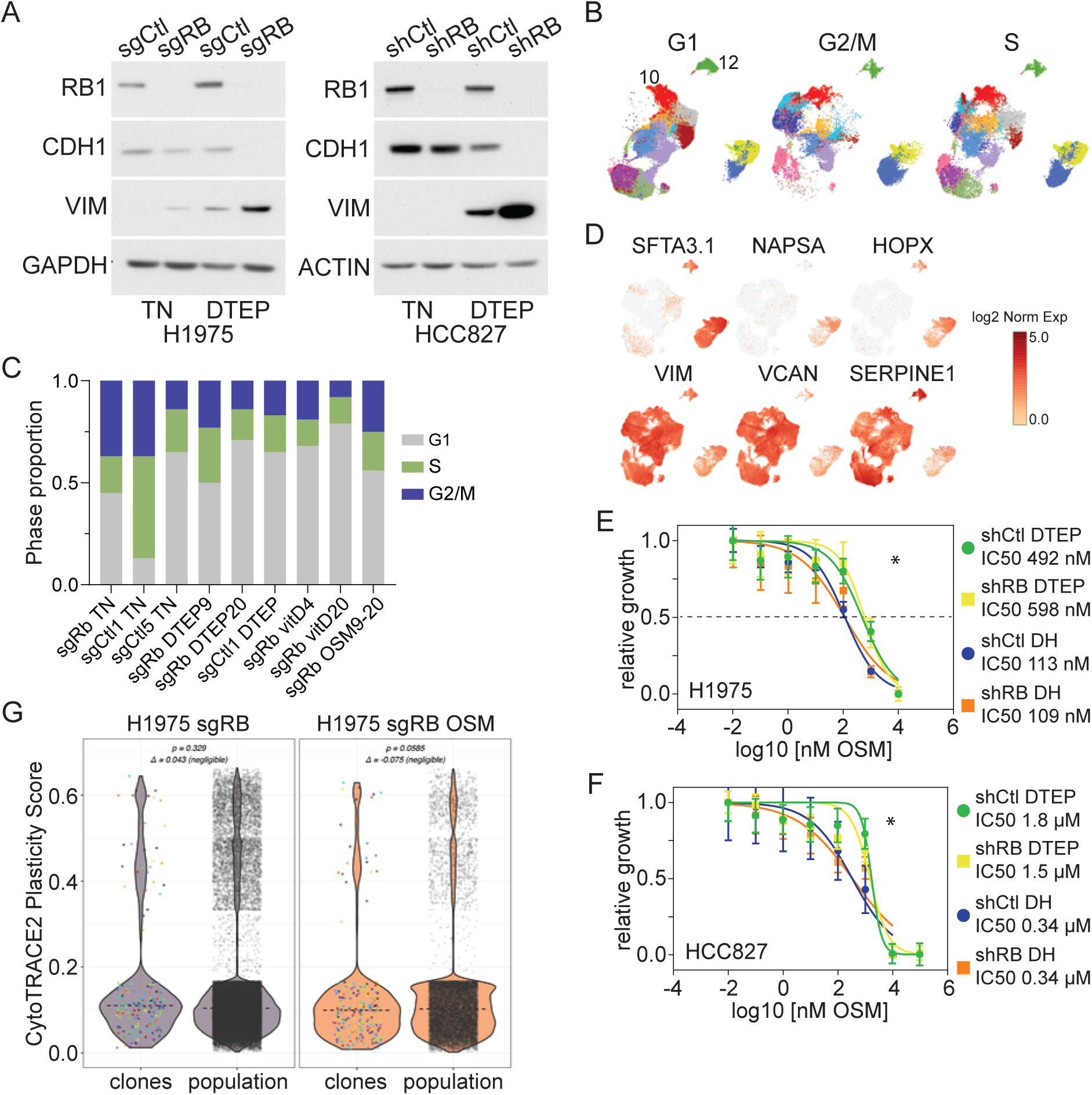
RB1 deficiency drives transcriptional plasticity enabling acquisition of reversible, OSM resistant transcriptional states. A) Protein extracted from the indicated H1975 or HCC827 cells was analyzed for epithelial (CDH1) or mesenchymal (VIM) marker protein expression by western blotting. GAPDH or ACTIN serve as protein loading controls. B) UMAPs of the scRNA-seq data in Figure 5D was segregated by cell cycle phase demonstrating the proliferation cells with OSM resistance associated gene expression clusters. C) scRNA-seq data was used to calculate the cell cycle phase distribution of cells within each sample, displayed in stacked columns. D) scRNA-seq UMAPs color coded based on log2 normalized expression of alveolar epithelial (*SFTA3.1, NAPSA, HOPX*) or mesenchymal (*VIM, VCAN, SERPINE1*) marker genes. E) The indicated H1975 OSM DTEP, either maintained on OSM (DTEP) or cultured in the absence of OSM during a drug holiday (DH), were tested for OSM sensitivity using 3-day dose response experiments. The non-linear least squares fit of the data from 3 biological replicates is plotted with best-fit IC50 concentrations shown. Asterisks indicate that the dose response curves were significantly different (P<0.05). F) The indicated HCC827 OSM DTEP, either maintained on OSM (DTEP) or cultured in the absence of OSM during a drug holiday (DH), were analyzed as in E). G) CytoTrace2 scores were calculated for cells within individual cell clones or the overall cell population and the distribution of scores displayed in violin plots. The distribution of scores for cell clones was not significantly different from the overall population (*P*>0.05).

**Figure S6:**
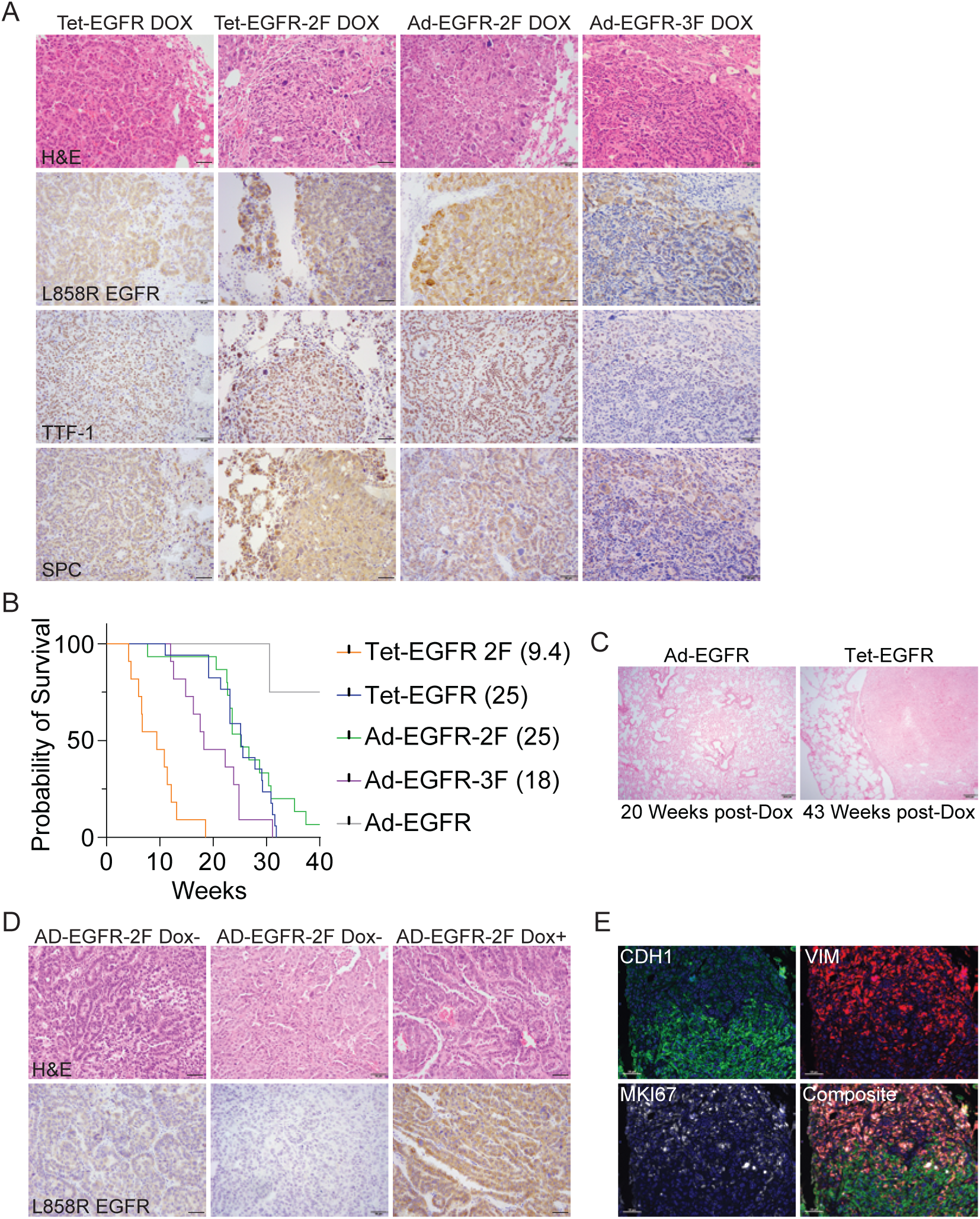
Tumor suppressor gene loss drives progression to mutant EGFR independent tumors. A) Representative images of lung tissue sections from mice of the indicated genotype maintained on Dox stained with H&E or immunostained for the indicated proteins. B) Overall survival of mice of the indicated genotypes maintained on Dox is shown on a Kaplan-Meier plot with median survival in weeks indicated in parentheses. Survival curves are significantly different based on log rank test (*P*<0.05). C) Lung tissue sections from an Ad-EGFR or Tet-EGFR mouse 20- or 43-weeks post-Dox withdrawal, respectively, were stained for H&E. D) Lung tissue sections from AdEGFR-2F mice either maintained on Dox or collected after relapse post-Dox withdrawal were stained with H&E or immunostained for L858R EGFR. Two examples of post-Dox lung tumors are shown with low or undetectable L858R EGFR expression. E) A lung tumor section from a mouse relapsing after Dox withdrawal was immunostained for the epithelial protein CDH1, the mesenchymal protein VIM, or the proliferation marker MKI67. The scale bars for all images in the figure represent 50 µM.

**Figure S7:**
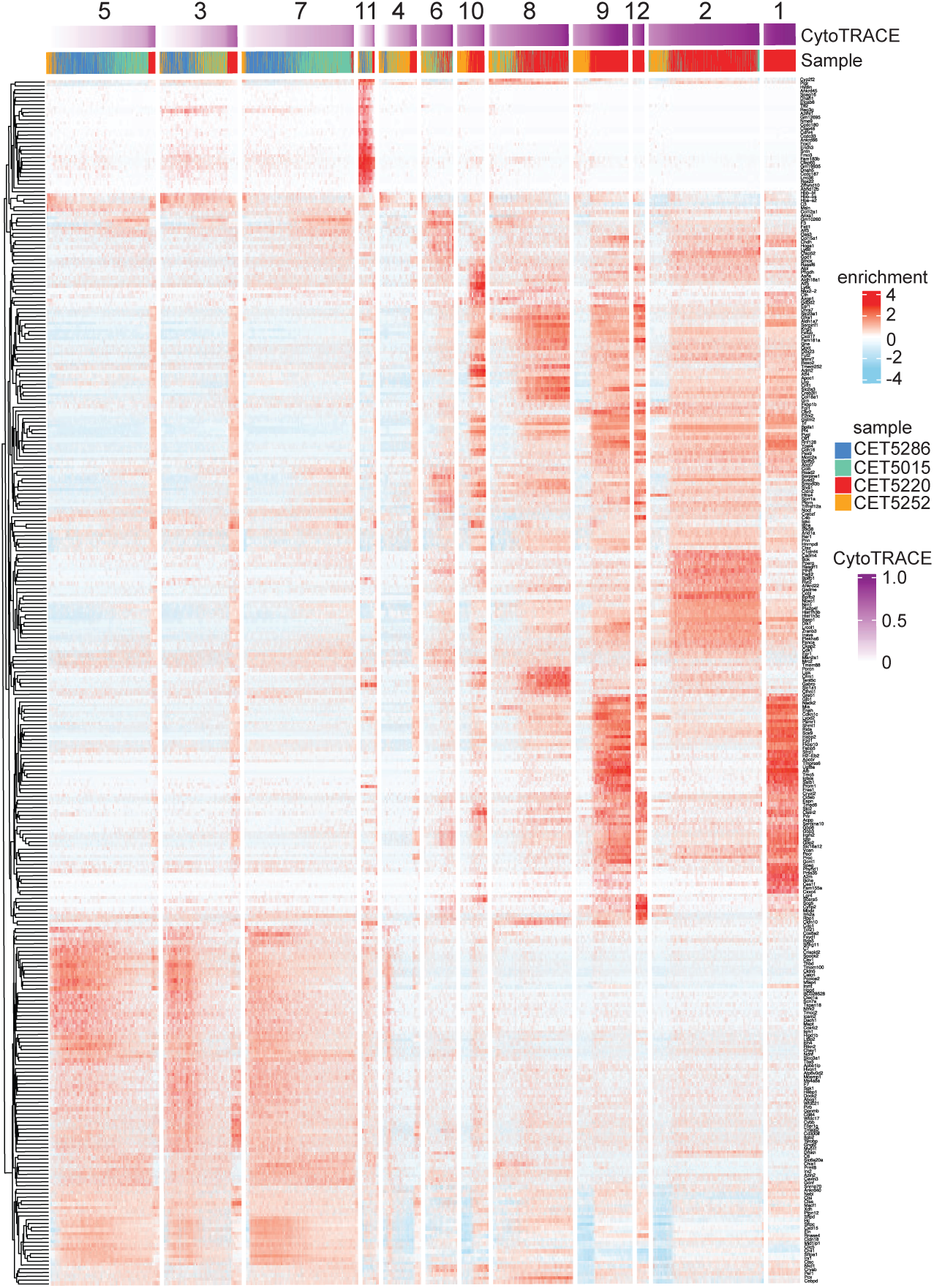
Heatmap of gene expression distinguishing transcriptional clusters detected in spatial transcriptomic profiling data. Spatial transcriptomic profiling identified 12 distinguishable transcriptional clusters across tissue sections from four mice with different treatments and genotypes. The heatmap identifies the most variable genes whose enrichment characterizes each of the 12 transcriptional clusters. Rows are composed of genes with columns composed of individual spots transcriptionally profiled. Sample identity and CytoTrace2 scores are at top.

**Figure S8:**
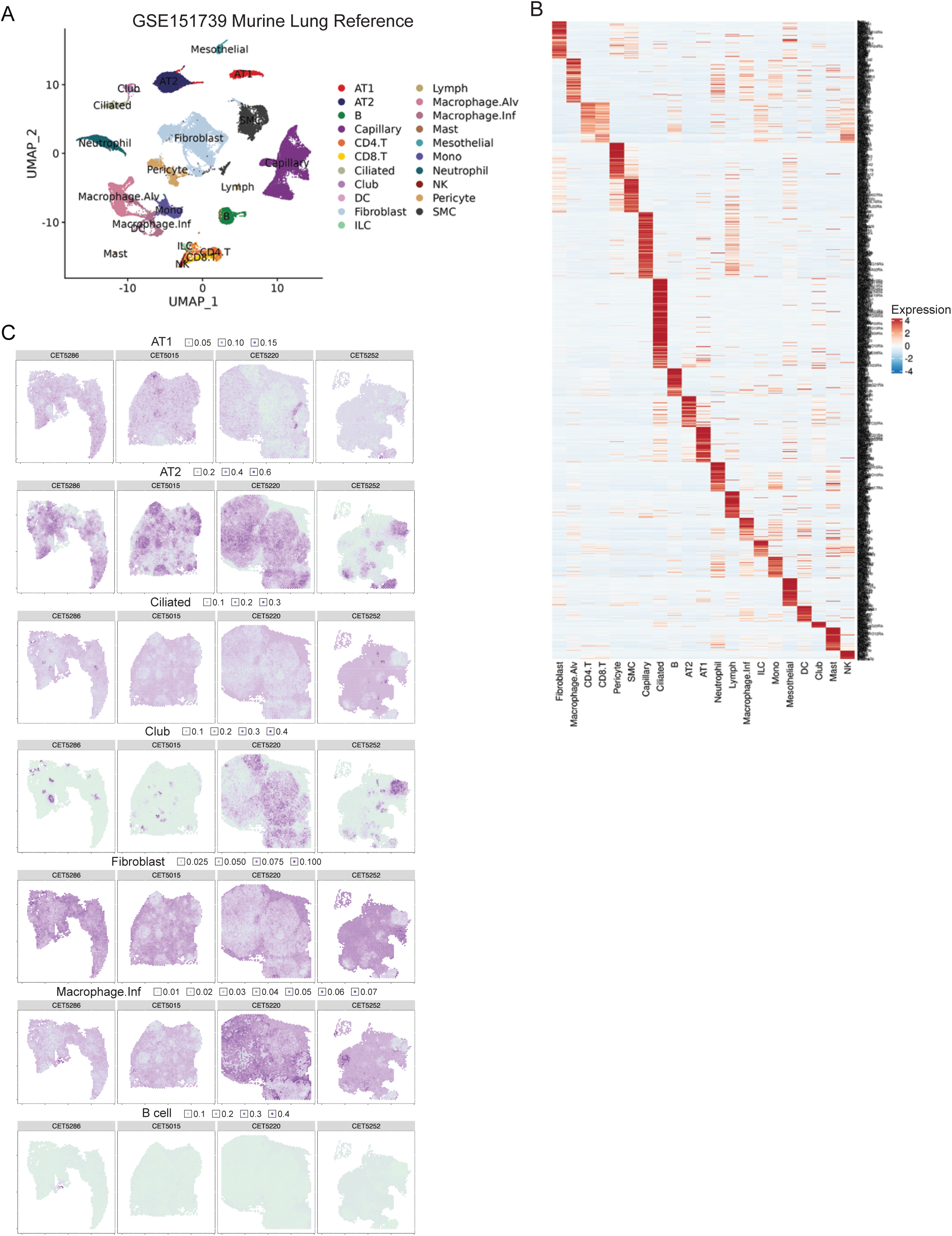
Reference mapping transcriptionally profiled lung tissue sections based on normal lung cell types. A) The indicated reference single cell RNA-seq data from normal mouse lung was clustered and displayed as a UMAP with each cluster identified by cell type. B) A heatmap showing variably expressed genes enriched within each identified cluster/cell type. The most enriched genes shown comprise the gene signature used to identify the corresponding cell type in the spatial transcriptomic data. C) Sections are shown with transcriptionally profiled spots color coded based on the signature score for the listed cell type.

