## Supplementary Table 7 for "Reversible therapeutic resistance in *EGFR* mutant lung cancer caused by *RB1* loss induced lineage plasticity"

| **Table S7: Top over-expressed genes in sgRb DTEP compared to sgRb TN (Log2 FC≥2.0)** | | | |
| --- | --- | --- | --- |
| **Gene** | **Expression is associated with** | | **References** |
| *ANKRD1* | EGFR TKI resistance, EMT | | (Ohara et al., 2021; Takahashi et al., 2018) |
| *PLCB4* | EGFR TKI resistance | | (Mao et al., 2021) |
| *TGFB2* | EGFR TKI resistance, EMT | | (Yang et al., 2022; Zhu et al., 2019) |
| *CEMIP* | EGFR signaling, EMT | | (Domanegg et al., 2022; Liu et al., 2023b) |
| *GPRC5A* | EGFR signaling, EMT | | (Lin et al., 2014; Zhang et al., 2023; Zhong et al., 2015) |
| *FGF13* | EGFR TKI resistance, EMT | | (Miyagi-Shiohira et al., 2021; Szymczyk et al., 2021) |
| *MB21D2* | EMT | | (Gracilla et al., 2020) |
| *CCN1* | EGFR TKI resistance, | | (Haque et al., 2020; Saab et al., 2019; Zhou et al., 2020) |
| *DDAH1* | EGFR TKI resistance, EMT | | (Huang et al., 2020; Ye et al., 2017) |
| *Col12A1* | EMT | | (Liu et al., 2023a; Tang et al., 2023) |
| *TENM3* |  | |  |
| *PCLO* | EGFR TKI resistance, | | (Frattini et al., 2017) |
| *NREP* | EMT | | (Liu et al., 2021) |
| *SLC7A5* | EGFR TKI resistance, EMT | | (Liu et al., 2022; Yoo and Han, 2022) |
| *CYP1B1* | EGFR TKI resistance, | | (Alfieri et al., 2011; Kwon et al., 2016; Zhang et al., 2022) |
| *FSTL1* | EMT | | (Liu et al., 2017; Sisto et al., 2022) |
| *TGFBR1* | EGFR TKI resistance, EMT | | (Zhang et al., 2021; Zhang et al., 2019) |

Alfieri, R. R., Galetti, M., Tramonti, S., Andreoli, R., Mozzoni, P., Cavazzoni, A., Bonelli, M., Fumarola, C., La Monica, S., Galvani, E.*, et al.* (2011). Metabolism of the EGFR tyrosin kinase inhibitor gefitinib by cytochrome P450 1A1 enzyme in EGFR-wild type non small cell lung cancer cell lines. Molecular cancer *10*, 143.

Domanegg, K., Sleeman, J. P., and Schmaus, A. (2022). CEMIP, a Promising Biomarker That Promotes the Progression and Metastasis of Colorectal and Other Types of Cancer. Cancers (Basel) *14*.

Frattini, M., Molinari, F., and Epistolio, S. (2017). The role of Piccolo in cancer treatment: relationship with EGFR and related therapies, and a marker for new targeted therapies. J Thorac Dis *9*, 4240-4243.

Gracilla, D. E., Korla, P. K., Lai, M. T., Chiang, A. J., Liou, W. S., and Sheu, J. J. (2020). Overexpression of wild type or a Q311E mutant MB21D2 promotes a pro-oncogenic phenotype in HNSCC. Mol Oncol *14*, 3065-3082.

Haque, I., Kawsar, H. I., Motes, H., Sharma, M., Banerjee, S., Banerjee, S. K., Godwin, A. K., and Huang, C. H. (2020). Downregulation of miR-506-3p Facilitates EGFR-TKI Resistance through Induction of Sonic Hedgehog Signaling in Non-Small-Cell Lung Cancer Cell Lines. Int J Mol Sci *21*.

Huang, Z., Ma, Y., Zhang, P., Si, J., Xiong, Y., and Yang, Y. (2020). Long non-coding RNA H19 confers resistance to gefitinib via miR-148b-3p/DDAH1 axis in lung adenocarcinoma. Anticancer Drugs *31*, 44-54.

Kwon, Y. J., Baek, H. S., Ye, D. J., Shin, S., Kim, D., and Chun, Y. J. (2016). CYP1B1 Enhances Cell Proliferation and Metastasis through Induction of EMT and Activation of Wnt/beta-Catenin Signaling via Sp1 Upregulation. PLoS ONE *11*, e0151598.

Lin, X., Zhong, S., Ye, X., Liao, Y., Yao, F., Yang, X., Sun, B., Zhang, J., Li, Q., Gao, Y.*, et al.* (2014). EGFR phosphorylates and inhibits lung tumor suppressor GPRC5A in lung cancer. Molecular cancer *13*, 233.

Liu, S., Chen, L., Zeng, J., and Chen, Y. (2023a). A prognostic model based on the COL1A1-network in gastric cancer. Am J Transl Res *15*, 1640-1653.

Liu, T., Liu, Y., Miller, M., Cao, L., Zhao, J., Wu, J., Wang, J., Liu, L., Li, S., Zou, M.*, et al.* (2017). Autophagy plays a role in FSTL1-induced epithelial mesenchymal transition and airway remodeling in asthma. Am J Physiol Lung Cell Mol Physiol *313*, L27-L40.

Liu, Y., Hu, G., Li, Y., Kong, X., Yang, K., Li, Z., Lao, W., Li, J., Zhong, J., Zhang, S.*, et al.* (2023b). Research on the biological mechanism and potential application of CEMIP. Front Immunol *14*, 1222425.

Liu, Y., Ma, G., Liu, J., Zheng, H., Huang, G., Song, Q., Pang, Z., and Du, J. (2022). SLC7A5 is a lung adenocarcinoma-specific prognostic biomarker and participates in forming immunosuppressive tumor microenvironment. Heliyon *8*, e10866.

Liu, Y. J., Zeng, S. H., Hu, Y. D., Zhang, Y. H., and Li, J. P. (2021). Overexpression of NREP Promotes Migration and Invasion in Gastric Cancer Through Facilitating Epithelial-Mesenchymal Transition. Front Cell Dev Biol *9*, 746194.

Mao, K., Lin, F., Zhang, Y., and Zhou, H. (2021). Identification of Key Genes and Pathways in Gefitinib-Resistant Lung Adenocarcinoma using Bioinformatics Analysis. Evol Bioinform Online *17*, 11769343211023767.

Miyagi-Shiohira, C., Saitoh, I., Watanabe, M., and Noguchi, H. (2021). Gene Expression in Pancreatic Cancer-Like Cells and Induced Pancreatic Stem Cells Generated by Transient Overexpression of Reprogramming Factors. J Clin Med *10*.

Ohara, S., Suda, K., and Mitsudomi, T. (2021). Cell Line Models for Acquired Resistance to First-Line Osimertinib in Lung Cancers-Applications and Limitations. Cells *10*.

Saab, S., Chang, O. S., Nagaoka, K., Hung, M. C., and Yamaguchi, H. (2019). The potential role of YAP in Axl-mediated resistance to EGFR tyrosine kinase inhibitors. Am J Cancer Res *9*, 2719-2729.

Sisto, M., Ribatti, D., Ingravallo, G., and Lisi, S. (2022). The Expression of Follistatin-like 1 Protein Is Associated with the Activation of the EMT Program in Sjogren's Syndrome. J Clin Med *11*.

Szymczyk, J., Sluzalska, K. D., Materla, I., Opalinski, L., Otlewski, J., and Zakrzewska, M. (2021). FGF/FGFR-Dependent Molecular Mechanisms Underlying Anti-Cancer Drug Resistance. Cancers (Basel) *13*.

Takahashi, A., Seike, M., Chiba, M., Takahashi, S., Nakamichi, S., Matsumoto, M., Takeuchi, S., Minegishi, Y., Noro, R., Kunugi, S.*, et al.* (2018). Ankyrin Repeat Domain 1 Overexpression is Associated with Common Resistance to Afatinib and Osimertinib in EGFR-mutant Lung Cancer. Sci Rep *8*, 14896.

Tang, Z., Yang, Y., Zhang, Q., and Liang, T. (2023). Epigenetic dysregulation-mediated COL12A1 upregulation predicts worse outcome in intrahepatic cholangiocarcinoma patients. Clin Epigenetics *15*, 13.

Yang, Y., Li, S., Wang, Y., Zhao, Y., and Li, Q. (2022). Protein tyrosine kinase inhibitor resistance in malignant tumors: molecular mechanisms and future perspective. Signal Transduct Target Ther *7*, 329.

Ye, J., Xu, J., Li, Y., Huang, Q., Huang, J., Wang, J., Zhong, W., Lin, X., Chen, W., and Lin, X. (2017). DDAH1 mediates gastric cancer cell invasion and metastasis via Wnt/beta-catenin signaling pathway. Mol Oncol *11*, 1208-1224.

Yoo, H. C., and Han, J. M. (2022). Amino Acid Metabolism in Cancer Drug Resistance. Cells *11*.

Zhang, L., Yang, W., Yang, J., and Sun, F. (2023). GPRC5A regulates proliferation and oxidative stress by inhibiting the STAT3/Socs3/c-MYC pathway in hepatocellular carcinoma. J Clin Biochem Nutr *73*, 43-51.

Zhang, M., Zhang, Y. Y., Chen, Y., Wang, J., Wang, Q., and Lu, H. (2021). TGF-beta Signaling and Resistance to Cancer Therapy. Front Cell Dev Biol *9*, 786728.

Zhang, W., Ke, Y., Liu, X., Jin, M., and Huang, G. (2022). Drug resistance in NSCLC is associated with tumor micro-environment. Reprod Biol *22*, 100680.

Zhang, Y., Zeng, Y., Liu, T., Du, W., Zhu, J., Liu, Z., and Huang, J. A. (2019). The canonical TGF-beta/Smad signalling pathway is involved in PD-L1-induced primary resistance to EGFR-TKIs in EGFR-mutant non-small-cell lung cancer. Respir Res *20*, 164.

Zhong, S., Yin, H., Liao, Y., Yao, F., Li, Q., Zhang, J., Jiao, H., Zhao, Y., Xu, D., Liu, S.*, et al.* (2015). Lung Tumor Suppressor GPRC5A Binds EGFR and Restrains Its Effector Signaling. Cancer Res *75*, 1801-1814.

Zhou, Y., Li, H., Liang, X., Du, H., Suo, Y., Chen, H., Liu, W., Duan, R., Huang, X., and Li, Q. (2020). The CCN1 (CYR61) protein promotes skin growth by enhancing epithelial-mesenchymal transition during skin expansion. Journal of cellular and molecular medicine *24*, 1460-1473.

Zhu, X., Chen, L., Liu, L., and Niu, X. (2019). EMT-Mediated Acquired EGFR-TKI Resistance in NSCLC: Mechanisms and Strategies. Front Oncol *9*, 1044.
